# Serotonergic Neurons Mediate Operant Conditioning in *Drosophila* Larvae

**DOI:** 10.1101/2021.06.14.448341

**Authors:** Kristina T Klein, Elise C Croteau-Chonka, Lakshmi Narayan, Michael Winding, Jean-Baptiste Masson, Marta Zlatic

## Abstract

Observed across species, operant conditioning facilitates learned associations between behaviours and outcomes, biasing future action selection to maximise reward and avoid punishment. To elucidate the underlying neural mechanisms, we built a high-throughput tracker for *Drosophila melanogaster* larvae, combining real-time behaviour detection with closed-loop optogenetic and thermogenetic stimulation capabilities. We demonstrate operant conditioning in *Drosophila* larvae by inducing a bend direction preference through optogenetic activation of reward-encoding serotonergic neurons. Specifically, we establish that the ventral nerve cord is necessary for this memory formation. Our results extend the role of serotonergic neurons for learning in insects as well as the existence of learning circuits outside the mushroom body. This work supports future studies on the function of serotonin and the mechanisms underlying operant conditioning at both circuit and cellular levels.

## Introduction

Animals must rapidly alter their behaviour in response to environmental changes. An important adaptation strategy is associative learning (***Dickinson, 1981***; ***Rescorla, 1988***), in which an animal learns to predict an unconditioned stimulus (US) by the occurrence of a conditioned stimulus (CS). The US is often a punishing or rewarding event such as pain or the discovery of a new food source (***Pavlov, 1927***). The nature of the CS distinguishes two major associative learning types: classical conditioning (***Pavlov, 1927***) and operant conditioning (***Skinner, 1938***; ***Thorndike, 1911***).

In classical conditioning, the CS is an inherently neutral environmental stimulus such as a sound, odour, or visual cue. Pairing with an appetitive or aversive US leads to learned approach or avoidance of the CS in the future. Many vertebrates (***Andreatta and Pauli, 2015***; ***Brown et al., 1951***; ***Jones et al., 2005***; ***Braubach et al., 2009***) and invertebrates (***Takeda, 1961***; ***Vinauger et al., 2014***; ***Alexander et al., 1984***; ***Wen et al., 1997***; ***Scherer et al., 2003***; ***Davis, 2005***; ***Cognigni et al., 2018***; ***Vogt et al., 2014***) can make these associations. Across the animal kingdom, neural circuits have been identified as convergence sites for the external CS and the rewarding or punishing US (***Heisenberg et al., 1985***; ***Hawkins and Byrne, 2015***; ***Owald and Waddell, 2015***; ***Gründemann and Lüthi, 2015***; ***Caroni, 2015***; ***Tonegawa et al., 2015***). In classical conditioning of both larval and adult *Drosophila*, the mushroom body (MB) brain area serves this purpose (*Cognigni et al., 2018*; *Heisenberg et al., 1985*; *Heisenberg, 2003*; *Rohwedder et al., 2016*; *Vogt et al., 2014*; *Saumweber et al., 2018*; *Owald and Waddell, 2015*). In each larval brain hemisphere, the CS is encoded by a subset of the approximately 110 Kenyon cells (KCs) (*Aso et al., 2014a*; *Honegger et al., 2011*; *Berck et al., 2016*; *Lin et al., 2014*; *Owald and Waddell, 2015*; *Campbell et al., 2013*; *Turner et al., 2008*; *Eichler et al., 2017*), which synapse onto 24 MB output neurons (MBONs) driving approach or avoidance (*Aso et al., 2014b*; *Owald et al., 2015*; *Perisse et al., 2016*; *Séjourné et al., 2011*; *Saumweber et al., 2018*; *Shyu et al., 2017*; *Plaçais et al., 2013*; *Eichler et al., 2017*). KC to MBON connection strength is modulated by dopaminergic and octopaminergic neurons, which represent the rewarding or punishing US (*Schwaerzel et al., 2003*; *Schroll et al., 2006*; *Honjo and Furukubo-Tokunaga, 2009*; *Vogt et al., 2014*; *Saumweber et al., 2018*; *Waddell, 2013*). Activation of the MB-innervating PAM cluster dopaminergic neurons serves as both a necessary and sufficient reward signal in classical conditioning (*Rohwedder et al., 2016*; *Liu et al., 2012*; *Vogt et al., 2014*; *Waddell, 2013*; *Cognigni et al., 2018*).

In operant conditioning, the CS is an animal’s own action (***Skinner, 1938***; ***Thorndike, 1911***). After memory formation, the animal can predict the outcome of its behaviour and bias future action selection accordingly, usually to maximise reward and avoid punishment (***Skinner, 1938***). This behavioural adaptation can facilitate novel action sequences (***Topál et al., 2006***; ***Nottebohm, 1991***; ***Fee and Goldberg, 2011***) and, in some cases, repetitive, high-frequency motor activity (***Olds and Milner, 1954***; ***Corbett and Wise, 1980***; ***Jin and Costa, 2010***; ***Lovell et al., 2015***). Such observations have wider implications for understanding diseases including obsessive-compulsive disorder and addiction (***Everitt et al., 2018***; ***Balleine et al., 2015***; ***Joel, 2006***). Invertebrates are also capable of operant conditioning (***Brembs, 2003***; ***Hoyle, 1979***; ***Abramson et al., 2016***; ***Nuwal et al., 2012***; ***Booker and Quinn, 1981***). Despite countless operant conditioning experiments across species using various CS–US combinations, the underlying neural mechanisms remain poorly understood. For an animal to associate an action with its outcome, behavioural information must converge with circuits encoding positive or negative valence. Although vertebrate basal ganglia-like structures exemplify this (***Fee and Goldberg, 2011***; ***Redgrave et al., 2011***; ***Balleine et al., 2009***), some learned action-outcome associations do not require the brain (***Booker and Quinn, 1981***; ***Horridge, 1962***; ***Grau et al., 1998***). Operant conditioning may hence occur in more than one area of the central nervous system (CNS). It is also unclear to what extent learning at these sites is mediated by synaptic plasticity (***Lovinger, 2010***; ***Surmeier et al., 2007***; ***Reynolds and Wickens, 2002***; ***Joynes et al., 2004***; ***Gómez-Pinilla et al., 2007***) versus changes in the intrinsic excitability of individual neurons (***Dong et al., 2006***; ***Shen et al., 2005***; ***Nargeot et al., 1997***; ***Brembs et al., 2002***; ***Nargeot et al., 2009***). We aim to establish the *Drosophila* larva as a tractable model system for studying the neural circuit mechanisms underlying operant conditioning.

*Drosophila melanogaster* larvae perform various different actions. Typically, when exploring an environment, a larva alternates between crawling via forward peristalsis (***Heckscher et al., 2012***) and bending its head once or more to the left or right (***Gomez-Marin et al., 2011***; ***Luo et al., 2010***; ***Kane et al., 2013***; ***Figure 1A***). In the presence of nociceptive stimuli, larvae exhibit escape behaviour. While the most common response is an increase in bending away from undesirable conditions, including extreme temperature (***Luo et al., 2010***; ***Lahiri et al., 2011***), light (***Kane et al., 2013***), or wind (***Jovanic et al., 2019***), larvae also retreat from aversive sources using backward peristalsis (***Masson et al., 2020***; ***Kernan et al., 1994***; ***Heckscher et al., 2012***; ***Vogelstein et al., 2014***; ***Figure 1A***). The fastest escape response is rolling, where the larva moves laterally by curling into a C-shape and quickly turning around its own body axis (***Robertson et al., 2013***; ***Hwang et al., 2007***; ***Ohyama et al., 2013***; ***Figure 1A***). In nature, rolling is only observed after exposure to a strong noxious stimulus, such as heat or a predator attack (***Ohyama et al., 2015***; ***Robertson et al., 2013***; ***Tracey et al., 2003***).

**Figure 1.**
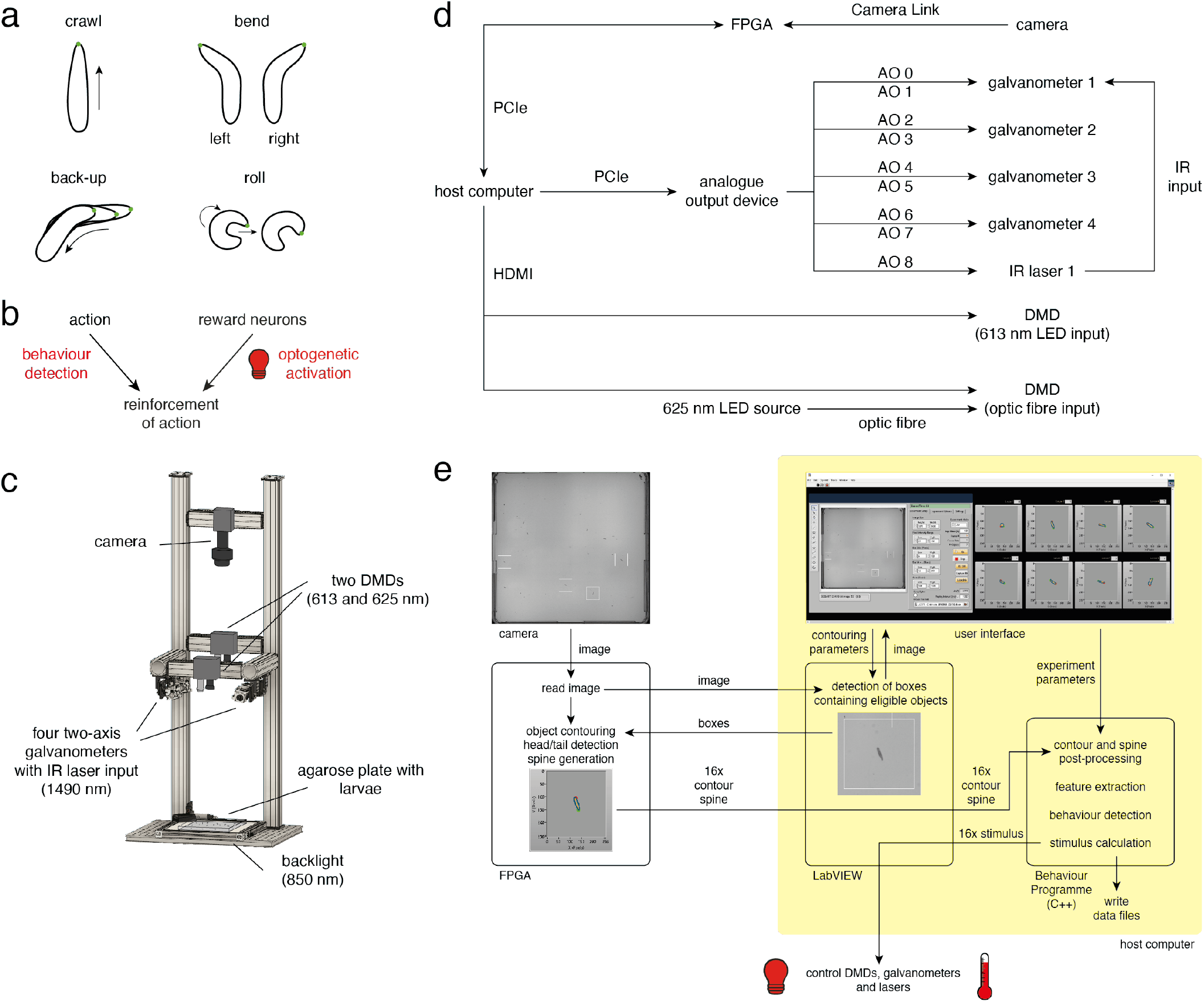
High-throughput operant conditioning in *Drosophila* larvae. **a.** Behavioural repertoire of *Drosophila* larvae. Schematics show the played by *Drosophila* larvae (crawl, left and right bend, backup and roll). The larval contour is displayed as a black outline with a green dot marking the head. **b.** In fully automated operant conditioning, an action of interest was reinforced by coupling real-time behaviour detection with optogenetic activation of reward circuits. **c.** High-throughput tracker schematic showing the relative positions of the agarose plate, backlight, camera, digital micromirror devices (DMDs), and galvanometers. IR: infrared. **d.** Block diagram of hardware components. AO: analogue output, FPGA: field-programmable gate array. **e.** Data flow between software elements. **Figure 1–Figure supplement 1.** Contour calculation on field-programmable gate array (FPGA). **Figure 1–Figure supplement 2.** Detecting head and tail. **Figure 1–Figure supplement 3.** Calculating a smooth spine and landmark points. **Figure 1–Figure supplement 4.** Calculating direction vectors. **Figure 1–Figure supplement 5.** Features describing body shape. **Figure 1–Figure supplement 6.** Velocity features. **Figure 1–Figure supplement 7.** Temporal smoothing of features. **Figure 1–Figure supplement 8.** Differentiation by convolution.

Powerful genetic toolkits have advanced the observation and manipulation of larval behaviour at the cellular level, making *Drosophila* larvae particularly well-suited for studying the neural mechanisms underlying learning. In *Drosophila*, individual neurons are uniquely identifiable, with morphology and function preserved across animals (***Skeath and Thor, 2003***; ***Wong et al., 2002***; ***Marin et al., 2002***; ***Jefferis et al., 2007***). Together with tissue-localised protein expression afforded by the GAL4-UAS binary expression system (***Fischer et al., 1988***; ***Brand and Perrimon, 1993***), this has yielded neuron-specific GAL4 drivers (***Jenett et al., 2012***; ***Luan et al., 2006***; ***Pfeiffer et al., 2010***) that reproducibly target the same group of cells in each individual. Adding fluorescent markers helps pinpoint a neuron’s location and reveal its anatomical features (***Lee and Luo, 1999***), while producing light-sensitive channelrhodopsins and temperature-sensitive ion channels facilitates optogenetic (***Zemelman et al., 2002***; ***Lima and Miesenböck, 2005***) or thermogenetic (***Hamada et al., 2008***; ***Kitamoto, 2001***) modulation of neural activity. Furthermore, the larva’s compact CNS has made it feasible to manually reconstruct neurons and their synaptic partners from a larval electron microscopy (EM) volume (***Berck et al., 2016***; ***Eichler et al., 2017***; ***Fushiki et al., 2016***; ***Ohyama et al., 2015***; ***Schlegel et al., 2016***; ***Larderet et al., 2017***; ***Jovanic et al., 2016, 2019***), giving rise to a full wiring diagram of the MB (***Eichler et al., 2017***; ***Eschbach et al., 2020a***,b).

There is overwhelming evidence that larvae are capable of classical conditioning. They can be trained to approach an odour paired with a gustatory reward (***Schleyer et al., 2011***; ***Hendel et al., 2005***; ***Kudow et al., 2017***; ***Niewalda et al., 2008***), or avoid an odour paired with light (***von Essen et al., 2011***), electric shock (***Aceves-Piña and Quinn, 1979***; ***Tully et al., 1994***), heat (***Khurana et al., 2012***), vibration (***Eschbach et al., 2011***), or the bitter compound quinine (***Gerber and Hendel, 2006***; ***Apostolopoulou et al., 2014***). Light can also be a CS: innate avoidance of light and preference for darkness (***Sawin-McCormack et al., 1995***) can be modulated when paired with reward or punishment (***Gerber et al., 2004***; ***von Essen et al., 2011***). It has remained an open question, however, whether *Drosophila* larvae can form action–outcome associations and where in the CNS these memories are formed.

Conducting operant conditioning with larvae requires real-time behaviour detection such that reward or punishment can be administered with minimal delay (***Figure 1B***). Single-animal closed-loop trackers have recently been developed (***Schulze et al., 2015***; ***Tadres and Louis, 2020***). However, the effciency of training paradigms would improve with automated US delivery and simultaneous conditioning of multiple animals. Therefore, we here introduce a high-throughput tracker for *Drosophila* larvae with real-time behaviour detection and closed-loop stimulation. Effciency of the setup stems from the simultaneous, real-time, behaviour detection for up to 16 freely moving larvae, and targeted closed-loop optogenetic and thermogenetic stimulus delivery with full intensity control and minimal delay.

## Results

### High-throughput closed-loop tracker

#### Hardware design

Designing an automated operant conditioning protocol for the *Drosophila* larva was challenging due to the larva’s physical characteristics. We excluded partial immobilisation protocols similar to the ones used to condition adult *Drosophila* navigation through virtual environments (***Nuwal et al., 2012***; ***Wolf and Heisenberg, 1991***; ***Wolf et al., 1998***; ***Brembs, 2011***). We instead built a high-throughput multi-larva tracker combining live computer vision behaviour detection with closed-loop control of US delivery in response to unrestricted larval behaviour.

All hardware resided within an optically opaque enclosure to ensure experiments were performed without environmental light. Larvae moved freely on an agarose plate, backlit from below by an infrared LED and observed from above through a high-resolution camera (***Figure 1C***). A Camera Link communication protocol interfaced with a high-performance field-programmable gate array (FPGA), which itself interacted with the host computer. The FPGA and the host computer performed image processing, behaviour detection, and stimulus calculation (***Figure 1D***).

Our operant conditioning paradigm targeted individual larvae performing specific behaviours. Optogenetic stimulation was achieved by directing red light through two digital micromirror devices (DMDs) which were programmed to project small 1 cm^2^ squares at the location of individual larvae. Both DMDs, which were positioned to project over the entire plate area, were operated simultaneously (***Figure 1C***).

Thermogenetic stimulation of individual larvae was achieved by directing a 1490 nm infrared (IR) laser beam through a two-axis scanning galvanometer mirror positioning system (***Figure 1C***), a technique previously used to stimulate single adult flies (***Bath et al., 2014***; ***Wu et al., 2014***). Because the 1490 nm wavelength is well-absorbed by water (***Curcio and Petty, 1951***), larvae exposed to the IR beam were rapidly heated. We took advantage of the galvanometer’s high scanning velocity to rapidly cycle the beam between four larvae (***Figure 1D***).

### Software architecture

Several computer vision algorithms exist for real-time tracking of freely behaving animals. ***Stowers et al. (2017)*** and ***Krynitsky et al. (2020)*** developed software for tracking mice, and ***Mischiati et al. (2015)*** developed high-speed tracking of single dragonflies in three-dimensional space. There are numerous tracking frameworks for adult *Drosophila*, some requiring the flies to move within a two-dimensional plane (***Straw and Dickinson, 2009***; ***Donelson et al., 2012***) while others detect the three-dimensional position of single (***Fry et al., 2008***) or multiple (***Grover et al., 2008***; ***Straw et al., 2011***) flies. The Multi-Worm Tracker (MWT) software developed by ***Swierczek et al.*** (***2011***) is suitable for simultaneously tracking a large number of *Caenorhabditis elegans* and has been adapted to analyse *Drosophila* larvae reactions in response to various stimuli (***Ohyama et al., 2013***; ***Vogelstein et al., 2014***; ***Jovanic et al., 2019***; ***Masson et al., 2020***).

Operant conditioning requires live behaviour detection to trigger delivery of reward or punishment. Numerous algorithms have been developed to analyse offline behavioural recordings of animals such as *C. elegans* (***Huang et al., 2006***; ***Stephens et al., 2008***; ***Gupta and Gomez-Marin, 2019***), zebrafish larvae (***Mirat et al., 2013***; ***Reddy et al., 2020***), adult *Drosophila* (***Katsov and Clandinin, 2008***; ***Branson et al., 2009***; ***Dankert et al., 2009***; ***Robie et al., 2017***; ***Berman et al., 2014***; ***Klibaite et al., 2017***), bees (***Veeraraghavan et al., 2008***), and mice (***Mathis et al., 2018***; ***Luxem et al., 2020***; ***van Dam et al., 2020***). The *Drosophila* larva has also attracted attention due to analytical challenges surrounding its deformable body and limited set of distinguishing features (***Luo et al., 2010***; ***Gomez-Marin et al., 2011***; ***Gershow et al., 2012***; ***Denisov et al., 2013***; ***Vogelstein et al., 2014***; ***Ohyama et al., 2013, 2015***; ***Masson et al., 2020***). Most of these approaches are not ideal to run in real time or require a mix of past and future information to provide reliable behaviour detection (***Gomez-Marin et al., 2011***; ***Masson et al., 2020***). More generally, machine learning based methods have gained momentum in providing both supervised and unsupervised approaches to behaviour analysis. It is worth noting a recent trend in developing unsupervised learning methods (e. g. ***Graving and Couzin, 2020***; ***Luxem et al., 2020***).

While real-time behaviour detection of casts and runs has been developed for a single animal (***Schulze et al., 2015***), our study of operant conditioning in freely behaving *Drosophila* larvae required effcient, real-time behaviour detection of multiple animals. We built a system to simultaneously track up to 16 larvae in real time, using LabVIEW for the user interface and algorithm implementation (***Figure 1E***). Instrumental to this software architecture was the fast image processing speed afforded by FPGA-based parallelisation (***Soares dos Santos and Ferreira, 2014***; ***Li et al., 2011***; ***Zhang et al., 2017***). Neuroscientists have adapted FPGA’s real-time analysis capabilities (***Shirvaikar and Bushnaq, 2009***; ***Uzun et al., 2005***; ***Chiuchisan, 2013***; ***Yasukawa et al., 2016***) to track rats (***Chen et al., 2005***), zebrafish larvae (***Cong et al., 2017***), and fluorescently labelled neurons in freely behaving *Drosophila* larvae (***Karagyozov et al., 2018***). In our system, the FPGA and host computer worked together to read the raw camera images, detect eligible objects, and extract and process object features (i. e. contour, head and tail position, and body axis) (***Figure 1E***). Larval body shape, velocity, and direction of motion facilitated robust behaviour detection which, in turn, drove closed-loop optogenetic and thermogenetic stimulation. All relevant experiment parameters and time-series data were output for offline analysis through a custom MATLAB framework (see Materials and methods).

Optogenetic and thermogenetic stimulation efficiency verified by behavioural readout We conducted proof-of-principle experiments to ensure that our set-up could be successfully used for optogenetic stimulation (***Figure 2A***). ***Ohyama et al.*** (***2015***) have identified two GAL4 lines expressed in neurons whose activation triggers strong rolling behaviour. *69F06-Gal4* drives expression in command neurons for rolling, whereas *72F11-Gal4* drives expression in the Basin neurons, which integrate mechanosensory and nociceptive stimuli. ***Klapoetke et al.*** (***2014***) have developed the red-shifted channelrhodopsin *CsChrimson*, which can be expressed under GAL4 control. We tested whether *69F06-Gal4 x UAS-CsChrimson* and *72F11-Gal4 x UAS-CsChrimson* larvae rolled upon exposure to red light (***Figure 2B***, see also Materials and methods). In each stimulation cycle, we observed above-threshold rolls in over 40% of *69F06-Gal4 x UAS-CsChrimson* larvae and over 70% of *72F11-Gal4 × UAS-CsChrimson* larvae. This behaviour significantly contrasted with that of *attP2 x UAS-CsChrimson* control larvae (***Figure 2C***), suggesting that the DMDs could be used for optogenetic stimulation without activating the animals’ photoreceptors.

**Figure 2.**
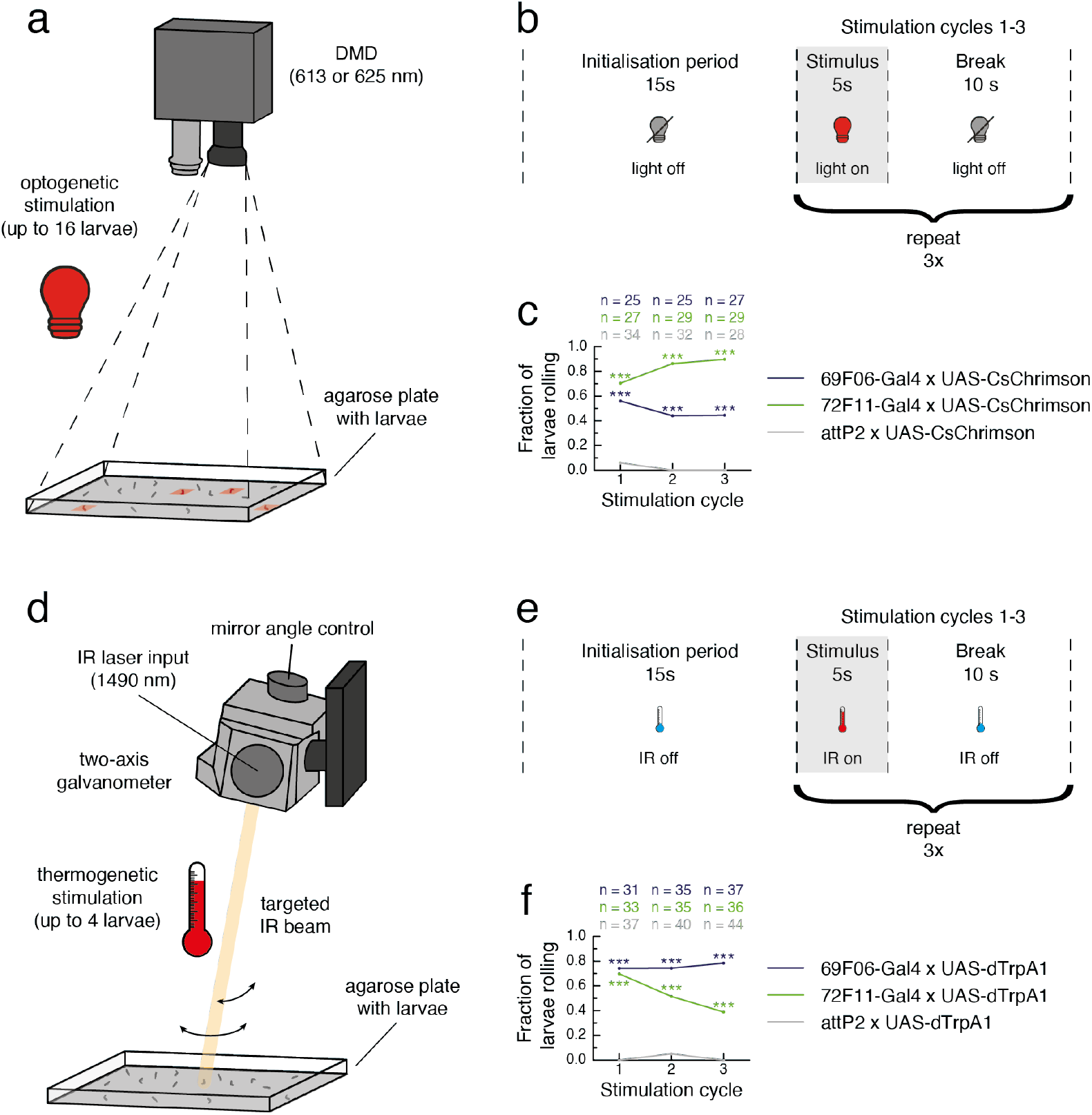
Optogenetic and thermogenetic stimulation with the high-throughput tracker. **a.** Hardware design schematic for optogenetic stimulation. Although the high-throughput tracker included two digital micromirror devices (DMDs), only one is shown for simplicity. **b.** Proof-of-principal experiment protocol for optogenetic stimulation. **c.** The fraction of larvae for which a roll was detected in each stimulation cycle. *69F06-Gal4 x UAS-CsChrimson* and *72F11-Gal4 x UAS-CsChrimson* larvae (*CsChrimson* expressed in neurons triggering roll behaviour; experiment groups) were compared to *attP2 x UAS-CsChrimson* larvae (no *CsChrimson* expression; control group). Fisher’s exact test was used to calculate statistical differences between the experiment and control groups (*** *p <* 0.001). **d.** Hardware design schematic for thermogenetic stimulation. Although the high-throughput tracker included four two-axis galvanometers, only one is shown for simplicity. IR: infrared. **e.** Proof-of-principal experiment protocol for thermogenetic stimulation. **f.** The fraction of larvae for which a roll was detected in each stimulation cycle. *69F06-Gal4 x UAS-dTrpA1* and *72F11-Gal4 x UAS-dTrpA1* larvae (*dTrpA1* expressed in neurons triggering roll behaviour; experiment groups) were compared to *attP2 x UAS-dTrpA1* larvae (no *dTrpA1* expression; control group). Fisher’s exact test was used to calculate statistical differences between the experiment and control groups (*** *p <* 0.001).

We also verified the efficacy of the galvanometer set-up for thermogenetic stimulation (***Figure 2D***). We tested whether *69F06-Gal4 x UAS-dTrpA1* and *72F11-Gal4 x UAS-dTrpA1* larvae rolled upon exposure to the IR laser (***Figure 2E***, see also Materials and methods). In each stimulation cycle, we observed above-threshold rolls in over 70% of *69F06-Gal4 x UAS-dTrpA1* larvae and over 35% of *72F11-Gal4 x UAS-dTrpA1* larvae; a significant contrast to the *attP2 x UAS-dTrpA1* control larvae whose roll rate was close to zero. We concluded that these heating conditions were effective for targeted *Trp* channel activation without larvae perceiving strong pain (***Figure 2F***).

### Operant conditioning of larval bend direction

We chose optogenetic activation of reward circuits as a US for automated operant conditioning. The main challenge was determining which neurons could convey a sufficient reinforcement signal, especially as the capacity for *Drosophila* larvae to exhibit operant learning was not yet demonstrated. Across the animal kingdom, it has been observed that biogenic amine neurotransmitters can provide such a signal (***Giurfa, 2006***; ***Hawkins and Byrne, 2015***; ***Meneses and Liy-Salmeron, 2012***; ***Fee and Goldberg, 2011***). It is also conceivable that the *Drosophila* PAM cluster dopaminergic neurons that can signal reward in classical conditioning (***Rohwedder et al., 2016***; ***Liu et al., 2012***; ***Vogt et al., 2014***; ***Cognigni et al., 2018***; ***Waddell, 2013***) may perform similarly in operant conditioning. We therefore aimed to induce operant conditioning by stimulating a broad set of dopaminergic and serotonergic neurons. If valence signalling relevant for operant conditioning is mediated by one of these two neurotransmitters, activation of this large set of neurons paired with behaviour should be sufficient to induce learning.

We expressed *UAS-CsChrimson* under the control of the *Ddc-Gal4* driver, which covers a large set of dopaminergic and serotonergic neurons in the CNS (***Li et al., 2000***; ***Sitaraman et al., 2008***; ***Lundell and Hirsh, 1994***), including the PAM cluster (***Liu et al., 2012***; ***Aso et al., 2012***). Although the function of most *Ddc* neurons is unknown, their collective activation can substitute for an olfactory conditioning reward in adult flies (***Liu et al., 2012***; ***Shyu et al., 2017***; ***Aso et al., 2012***). The goal of our paradigm was to establish a learned direction preference for bending, conditioning *Ddc-Gal4 x UAS-CsChrimson* larvae to bend more often to one side than the other. Although stimulation side was randomized across trials, we describe (for simplicity) the experiment procedure where this predefined side was the left. Each experiment began with a one-minute test period where no light was presented. What followed were four training sessions, each three-minutes long, in which larvae received optogenetic stimulation when bending to the left. Between training sessions, larvae experienced three minutes without stimulation. Larvae were periodically brushed back to the centre of the agarose plate to mitigate the experimental side effects of reaching the plate’s edge (see Materials and methods for more details). Following the fourth training session was a one-minute test period without stimulation (***Figure 3A***).

**Figure 3.**
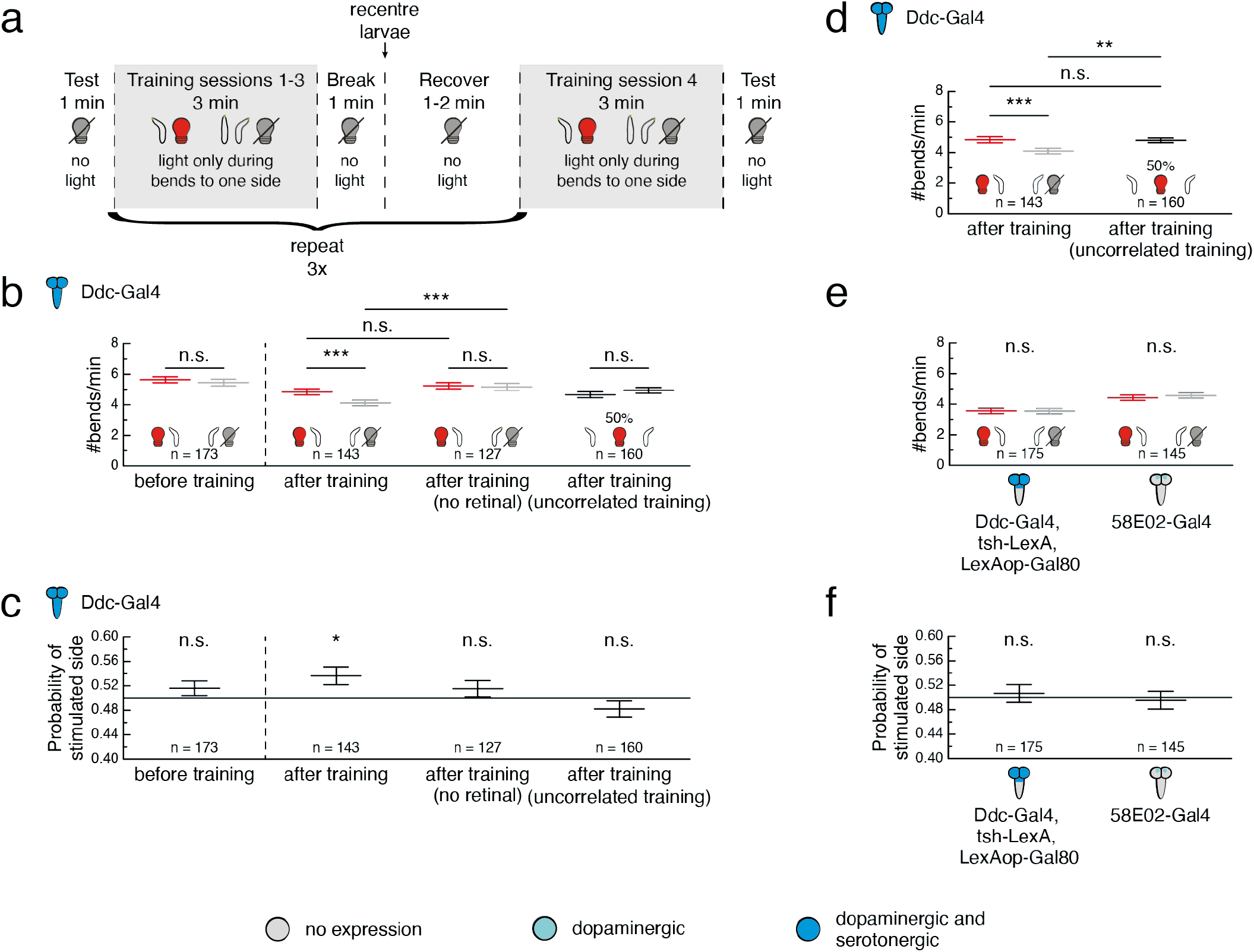
Operant conditioning of bend direction in *Drosophila* larvae requires the ventral nerve cord. **a.** Experiment protocol using the high-throughput closed-loop tracker. Behaviours are depicted as larval contours (black) with head (green). During training, the larva received an optogenetic stimulus (red light bulb) whenever it bent to one predefined side (here depicted as the left for simplicity), and light was switched off light bulb). **b,d,e.** Larval bend rate shown as the number of bends per minute, grouped by bend direction. The bend rate to the stimulated side (depicted as a left bend with a red light bulb for simplicity) is shown in red and the bend rate to the unstimulated side (depicted as a right bend with a grey light bulb for simplicity) is shown in grey. For larvae that received random, uncorrelated stimulation during 50% of bends, the bend rates to the left and right are shown in black. Statistical differences within groups were tested with a test; statistical differences between two groups were tested with a two-sided Wilcoxon signed-rank test; statistical differences between two groups were tested with a two-sided Mann-Whitney *U* test. **c,f.** Probability that a given bend was directed towards the stimulated side or, in the case of the uncorrelated training group, towards the left. Grey line indicates equal probability of 0.5 for bends to either side. Statistics calculated from a two-sided Wilcoxon signed-rank test. **b–f.** Gal4 expression depicted as color-coded CNS. All data is shown as (mean ± s. e. m.). n. s. *p* ≥ 0.05 (not significant), * *p <* 0.05, ** *p <* 0.01, *** *p <* 0.001. **b.** Bend rate for *Ddc-Gal4 x UAS-CsChrimson* larvae. Data is shown from the test period before the first training session and the test period after the fourth training session. **c.** Data from same experiments as in **b**. **d.** Same data as in **b**, but bend rate for uncorrelated training group was calculated without stratification by bend direction. **e.** Bend rate for *Ddc-Gal4 x UAS-CsChrimson; tsh-LexA, LexAop-Gal80* and *58E02-Gal4 x UAS-CsChrimson* larvae. The effector, *UAS-CsChrimson*, is omitted from the figure for visual clarity. Data is shown from the test period immediately following the fourth training session. **f.** Data from same experiments as in **e. Figure 3–Figure supplement 1.** *Ddc-Gal4* expression pattern without and with *tsh-Gal80* restriction.

For each larva, two measures served as a read-out for bend direction preference: i) the bend rate, measured as the number of bends per minute performed towards a given side, and ii) the probability that a given bend was directed towards the stimulated side, obtained by normalising the bend rate with the total number of bends performed by the larva in that minute. Individual larva variation in bend rate yielded different results for these measures at the population level. In the one-minute test prior to the first training session, we observed no significant difference in larval bend rate to either side and the likelihood of these naïve animals choosing one side over the other was not significantly different from chance. In the one-minute test following the fourth training session, larvae showed a preference for bends towards the side paired with red light stimulation during training, and the probability of these larvae bending towards this previously stimulated side was significantly greater than 50% (***Figure 3B***).

The light-dependent activation of neurons using *CsChrimson* requires a cofactor, retinal, which we supplemented in the food during development (***Klapoetke et al., 2014***; see Materials and methods). A control group of larvae raised on food without retinal showed no significant difference in absolute bend rate (***Figure 3B***) or bend direction probability (***Figure 3C***) throughout the experiment. This suggested that the US, which triggered a learned direction preference for bends in larvae raised on retinal, was indeed the collective activation of all *Ddc* neurons and not the red light itself. Notably, when directly comparing larvae raised with retinal to this control group raised without, the two groups showed no significant difference in the bend rate towards the stimulated side. Instead, the bend rate towards the unstimulated side was significantly reduced in larvae that received paired training compared to this control (***Figure 3B***). This raised the question whether larvae were learning to prefer the side paired with the rewarding US, or rather to avoid the side without the stimulus.

To confirm that the bend preference we observed after training was attributable to pairing light with bends solely in one direction, we conducted another control experiment in which larvae received random, uncorrelated stimulation during 50% of bends regardless of direction. After training, larvae showed neither a difference in absolute left and right bend rates, nor a significant probability of choosing one side over the other (***Figure 3B, Figure 3C***). These bend rates averaged together were indistinguishable from those of pair-trained larvae as they bent to the previously stimulated side. However, larvae which received uncorrelated training showed a significantly higher bend rate overall compared to pair-trained larvae bending to the previously unstimulated side (***Figure 3D***).

### The mushroom body is not sufficient to mediate operant conditioning in larvae

Our experiments showed that activation of *Ddc* neurons is a sufficient US for operant conditioning. While we did not identify which individual neurons mediate the observed effect, we hypothesised that not all *Ddc* neurons are involved. Some prior work in adult flies suggests that the MB is involved in operant conditioning (***Sun et al., 2020***), while other studies in the adult suggest that operant conditioning does not require the MB (***Booker and Quinn, 1981***; ***Wolf et al., 1998***; ***Colomb and Brembs, 2010, 2016***) and may instead involve motor neuron plasticity (***Colomb and Brembs, 2016***). The extent to which the MB is dispensable in larval operant conditioning is unknown. We investigated whether smaller subsets of *Ddc* neurons in the brain and subesophageal zone (SEZ) could support memory formation in our bend direction paradigm.

GAL80 under control of the *tsh* promoter suppresses GAL4 expression in the ventral nerve cord (VNC), but not in the brain or SEZ (***Clyne and Miesenböck, 2008***; ***Figure 3***–***Figure Supplement 1***). When trained under our operant conditioning protocol (***Figure 3A***), *Ddc-Gal4 x UAS-CsChrimson; tsh-LexA, LexAop-Gal80* larvae were equally likely to bend towards the side where they had previously received the optogenetic stimulus as they were to bend towards the unstimulated side (***Figure 3E, Figure 3F***). Activating these neurons was thus an insufficient rewarding US in this paradigm. The loss of the operant conditioning effect we observed with *Ddc-Gal4 x UAS-CsChrimson* larvae highlighted the necessity of dopaminergic or serotonergic neurons in the VNC for the formation of a bend direction preference. Their sufficiency was inconclusive, however, since perhaps two or more distinct groups of *Ddc* neurons needed collective activation in order to form a memory.

We then assessed whether exclusively activating the PAM cluster dopaminergic neurons innervating the MB could induce operant conditioning, as is the case for classical conditioning. *58E02-Gal4* drives expression in the majority of these neurons (***Rohwedder et al., 2016***). *58E02-Gal4 x UAS-CsChrimson* larvae did not develop any direction preference for bends following training (***Figure 3E, Figure 3F***). It is unsurprising that activation of these neurons alone could not act as a rewarding US in this paradigm, given our finding that *Ddc* neurons in the brain and SEZ are insufficient. It is remarkable, however, because it suggests that the neural circuits signalling reward in operant conditioning differ from those of classical conditioning. Although it remains to be seen whether these PAM cluster neurons contribute to memory formation by interacting with other *Ddc* neurons, these results further supported the idea that operant conditioning in *Drosophila* may not be mediated by the MB.

### Serotonergic neurons in brain and SEZ are a sufficient reward signal in classical conditioning

Pairing an action with activation of numerous dopaminergic and serotonergic neurons across the CNS was sufficient to induce operant conditioning of bend direction preference. Furthermore, our results indicated that the VNC subset of these neurons was essential for memory formation in the paradigm. It was an open question, however, whether this learning was mediated by dopamine, serotonin, or both. Dopamine and serotonin receptors are necessary for different classical conditioning tasks in honeybees, suggesting that the two neurotransmitters may carry out separate functions (***Wright et al., 2010***). We conducted a high-throughput classical conditioning screen of sparser dopaminergic and serotonergic driver lines to identify US candidates for comparison with our operant conditioning paradigm.

We expressed *CsChrimson* under the control of different GAL4 driver lines and tested whether pairing optogenetic activation of these neurons (US) with odour presentation (CS) could induce olfactory memory. Conditioning was performed using a similar procedure to those described in ***Gerber and Hendel*** (***2006***), ***Saumweber et al.*** (***2011***) and ***Eschbach et al.(2020b)***. In the paired group, larvae were exposed to alternating three-minute presentations of ethyl acetate with red light and air with no light. To ensure that any observed effects were a result of learning rather than innate odour preference or avoidance, an unpaired group was trained simultaneously with reciprocal stimulus presentation (odour/dark, air/light). Following training, larvae in both groups were tested on their preference for the odour in the absence of light (***Figure 4A***). All learning scores were compared to a negative control containing no GAL4 driver, *w*^*1118*^ × *UAS-CsChrimson*, which did not exhibit a learning phenotype (***Figure 4B***). Consistent with prior study results (***Rohwedder et al., 2016***; ***Eichler et al., 2017***; ***Almeida-Carvalho et al., 2017***), *58E02-Gal4 x UAS-CsChrimson* larvae showed appetitive olfactory learning with a significantly higher performance index than *w1118 x UAS-CsChrimson* larvae and so were used as a positive control (***Figure 4B***). *Ddc-Gal4 x UAS-CsChrimson* larvae exhibited appetitive memory comparable to *58E02-Gal4*(*p* = 0.1304, two-sided Mann-Whitney *U* test); an unsurprising result since the *Ddc-Gal4* expression pattern includes the PAM cluster neurons. Consistent with previous studies in the larva (***Schroll et al., 2006***) and adult (***Aso et al., 2012***; ***Claridge-Chang et al., 2009***; ***Liu et al., 2012***), *TH-Gal4 x UAS-CsChrimson* larvae exhibited significant aversive olfactory learning. *TH-Gal4* covers most dopaminergic neurons, excluding the PAM cluster (***Rohwedder et al., 2016***). The effect we observed may be mediated by punishment-signalling dopaminergic neurons that project to the MB vertical lobes (***Eschbach et al., 2020b***; ***Selcho et al., 2009***). Isolating the locus of this effect may prove challenging, given the dearth of larval driver lines targeting dopaminergic neurons without MB innervation.

**Figure 4.**
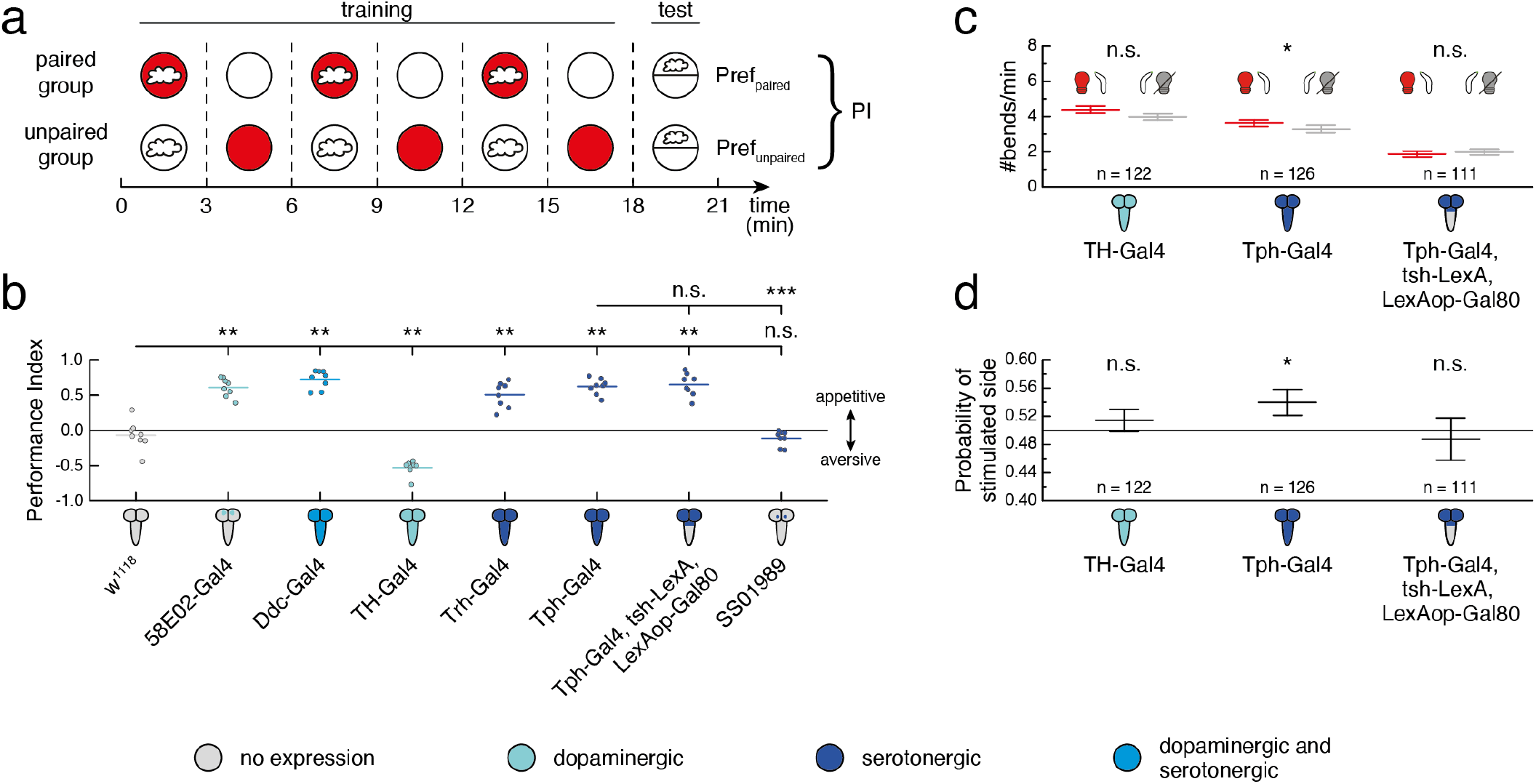
Different serotonergic neurons mediate classical and operant conditioning. All fly lines contained the *UAS-CsChrimson* effector, which is omitted from the figure for visual clarity. Gal4 expression depicted as color-coded CNS. **a.** Olfactory conditioning experiment protocol. During training, larvae in the paired group received three minutes of optogenetic red light stimulation (solid red circles) paired with the odour(white cloud) followed by three minutes of darkness (solid white circles) paired with air (no cloud). The unpaired group received reciprocal stimulus presentation (dark paired with odour, light paired with air). This procedure was repeated three times. In half of the experiments, the order of training trials was reversed, starting with air presentation instead of odour presentation. Both groups were then tested for learned odour preference in the dark with odour presented on one side of the plate and no odour on the other (PI = performance index). **b.** Performance indices following olfactory conditioning, plotted as raw data points and mean. *w*^*1118*^ × *UAS-CsChrimson* was the negative control (grey, *n* = 8), *58E02-Gal4 x UAS-Chrimson* was the positive control (blue, *n* = 8). Statistical comparisons to *w*^*1118*^ x *UAS-CsChrimson* were calculated using a two-sided Mann-Whitney *U* test with Bonferroni correction; n. s. *p ≥* 0.05/7 (not significant), ** *p <* 0.01/7. Statistical comparisons to *Tph-Gal4 x UAS-CsChrimson* were calculated using a two-sided Mann-Whitney *U* test with Bonferroni correction; n. s. *p* ≥ 0.05/2 (not significant), *** p < 0:001/2. **c,d.** All data is shown as (mean ± s. e. m.), n. s. *p ≥* 0.05 (not significant), * *p <* 0.05. **c.** Experiments followed the protocol depicted in ***Figure 3A***. Data is shown from the test period immediately following the fourth training session. Larval bend rate shown as the number of bends per minute, grouped by bend direction. The bend rate to the stimulated side (depicted as a left bend with a red light bulb for simplicity) is shown in red and the bend rate to the unstimulated side (depicted as a right bend with a grey light bulb for simplicity) is shown in grey. Statistical differences within a group were tested with a two-sided Wilcoxon signed-rank test. **d.** Probability that a given bend is directed towards the stimulated side. Grey line indicates equal probability of 0.5 for bends to either side. Statistics were based on a two-sided Wilcoxon signed-rank test. **Figure 4–Figure supplement 1.** *Tph-Gal4* expression pattern without and with *tsh-Gal80* restriction. **Figure 4–Figure supplement 2.** SS01989 exclusively drives expression in the CSD neuron. **Figure 4–Figure supplement 3.** Paired and unpaired group data for olfactory conditioning experiments.

Serotonergic signalling is required for associative learning in both larval (***Huser et al., 2017***) and adult (***Johnson et al., 2011***; ***Sitaraman et al., 2012***) *Drosophila*. We tested *Trh-Gal4* and *Tph-Gal4*, two driver lines that target the majority of serotonergic neurons and no dopaminergic neurons across the CNS of third-instar larvae (***Huser et al., 2012***). Consistent with previous reports (***Ganguly et al., 2020***), larvae expressing *CsChrimson* under either driver line formed strong appetitive olfactory memory, highlighting the sufficiency of serotonin as a US in associative learning. *Tph-Gal4* targets fewer serotonergic neurons than *Trh-Gal4*, making it valuable for narrowing down which serotonergic neurons serve as a relevant reward signal. We eliminated all *Tph-Gal4* expression in the VNC using *tsh-Gal80* (***Figure 4***–***Figure Supplement 1***). Activating the remaining *Tph* neurons in the brain and SEZ was sufficient to induce strong appetitive memory (***Figure 4B***). This result was notable and raised further questions: are serotonergic neurons in the brain and SEZ indirectly connected to MB-innervating dopaminergic neurons or do alternative learning circuits exist that altogether bypass the MB?

The contralaterally projecting serotonin-immunoreactive deutocerebral (CSD) neuron (***Roy et al., 2007***) is one previously described serotonergic brain neuron within the *Tph-Gal4* expression pattern (***Huser et al., 2012***) that innervates the antennal lobe and only has a few indirect connections to the MB (***Berck et al., 2016***). Combining anatomical features from existing EM reconstruction (***Berck et al., 2016***) with available lineage information facilitated identification of a split-GAL4 line (*SS01989*) that drives expression exclusively in the CSD neuron (***Figure 4***–***Figure Supplement 2***). Pairing activation of *SS01989* with ethyl acetate was insufficient for inducing olfactory memory (***Figure 4B***), suggesting that the classical conditioning phenotype we observed under *Tph-Gal4 x UAS-CsChrimson; tsh-LexA, LexAop-Gal80* was mediated by at least one other group of serotonergic neurons in the brain or SEZ.

### Serotonergic neurons in VNC are necessary for operant conditioning of bend direction

Given their strong associative learning phenotypes, we used the *TH-Gal4* and *Tph-Gal4* drivers to investigate whether operant conditioning of bend direction could be induced exclusively by dopaminergic or serotonergic neurons, respectively. Under our high-throughput training protocol (***Figure 3A***), *TH-Gal4 x UAS-CsChrimson* larvae showed no difference in bend rate between the previously stimulated and unstimulated sides in the one-minute test period (***Figure 4C***). Furthermore, the probability that any given bend was directed towards the previously stimulated side was not significantly different from chance (***Figure 4D***). Activating these dopaminergic neurons was an insufficient substitute for reward or punishment in operant conditioning.

Paired activation of *Tph-Gal4* neurons during bends to one side resulted in a significantly higher bend rate to the stimulated side relative to the unstimulated side during the test period (***Figure 4C***). The probability of bending in the previously stimulated direction was also significantly elevated (***Figure 4D***). In this way, activation of *Tph*-positive serotonergic neurons paired with bends to one side was sufficient for the formation of a learned direction preference. Combining this result with the knowledge that operant conditioning was impaired following restriction of *Ddc-Gal4 x UAS-CsChrimson* expression to the brain and SEZ suggests that the serotonergic neurons of the VNC were necessary for memory formation in this paradigm. Because *Tph-Gal4* is a broad driver line, it is possible that its expression pattern contains brain or SEZ neurons outside of those in *Ddc-Gal4*. The existence of these neurons could have potentially induced learning through an alternate mechanism independent from that which drove memory formation following *Ddc* neuron activation.

To assess whether the VNC serotonergic neurons were necessary for the observed operant conditioning effect, we used *tsh-Gal80* to restrict the *Tph-Gal4* expression pattern to the brain and SEZ. Paired optogenetic activation of *Tph-Gal4 x UAS-CsChrimson; tsh-LexA, LexAop-Gal80* with larval bends to one side was insufficient for operant conditioning (***Figure 4C, Figure 4D***). The *Tph-Gal4* expression pattern contains two neurons per VNC hemisegment (with the exception of a single neuron in each A8 abdominal hemisegment), all of which are serotonergic (***Huser et al., 2012***). While there are few serotonergic VNC candidates, we could not conclude from our data whether the operant conditioning effect relied solely on these neurons or whether synergistic activity from both the VNC and the brain or the SEZ was needed. Testing these hypotheses remains challenging since, to our knowledge, no sparse driver lines exist to exclusively target VNC serotonergic neurons.

Under a classical conditioning paradigm, we have confirmed that there exist learning pathways in *Drosophila* that rely on serotonergic neurons. We have also shown that serotonergic neurons can serve as a sufficient US for operant conditioning. Notably, different circuit mechanisms underlie classical and operant conditioning mediated by serotonergic neurons: activation of the brain and SEZ is sufficient for classical conditioning, whereas the VNC is necessary for operant conditioning.

## Discussion

Due to available genetic tools and the emerging connectome, the *Drosophila* larva is a uniquely advantageous model organism for neuroscience. We have uncovered a previously unknown neuronal mechanism of operant conditioning in the *Drosophila* larva. Serotonergic signalling can be employed as a reinforcing US in both classical and operant associative learning, but we are mindful that a single neural mechanism for learning may not exist. Distinct types of learning may share neurotransmitters or circuit components, but there may remain fundamental differences in connectivity and function. The experimental system we built was instrumental in investigating the neural circuits of operant conditioning, as it combined FPGA-based real-time tracking of multiple larvae with robust online behaviour detection and closed-loop stimulus presentation. This system could facilitate further research in taxis (***Luo et al., 2010***; ***Gomez-Marin et al., 2011***; ***Kane et al., 2013***; ***Jovanic et al., 2019***), decision-making (***Krajbich, 2019***; ***DasGupta et al., 2014***), and spatial navigation and memory (***Neuser et al., 2008***; ***Haberkern et al., 2019***). While further work is necessary, our bend direction paradigm provides a strong foundation for continued study of the circuit and cellular mechanisms underlying operant conditioning.

### High-throughput operant conditioning in Drosophila larvae

We have shown that *Drosophila* larvae are capable of operant conditioning and that optogenetic activation of *Ddc* neurons serves as a rewarding US during this learning process. With training, larvae formed an association between the presence or absence of this US and the direction in which they were bending. During testing, in the absence of any stimulation, larvae showed a significant learned preference for bending towards the previously stimulated side. This learned modification of future behaviour was only observed when the CS and US were paired during training; a hallmark of operant conditioning. Because *Ddc-Gal4* drives expression in dopaminergic and serotonergic neurons (***Li et al., 2000***; ***Sitaraman et al., 2008***), we concluded that one or both neurotransmitters are involved in memory formation under these experiment conditions.

Strong parallels exist between our operant learning paradigm and those employed for conditioning direction preference in adult *Drosophila*. Consider the work of ***Nuwal et al.*** (***2012***), in which tethered flies walked on a rotating ball and were rewarded with optogenetic activation of sugar-sensing neurons upon turns to one direction. As a consequence of this pairing, the flies learned to increase the number of turns to this side. Notably, the nature of the US remains an important difference between these paradigms. Our initial attempts to operantly condition larvae using activation of sugar-sensing neurons as a rewarding US were unsuccessful. These neurons, defined by two different *Gr43a-Gal4* drivers, were also insufficient for memory formation when activated with a paired odour in an olfactory conditioning screen. This was surprising, considering extensive evidence that natural sugar can serve as a rewarding US for classical conditioning in larvae (***Apostolopoulou et al., 2013***; ***Schleyer et al., 2015***; ***Weiglein et al., 2019***; ***Honjo and Furukubo-Tokunaga, 2005***; ***Neuser et al., 2005***; ***Rohwedder et al., 2012***; ***Scherer et al., 2003***; ***Schipanski et al., 2008***). One possible explanation for these discrepancies is that multiple groups of sensory neurons must be co-activated in order to relay a meaningful reward signal. Alternatively, it may be necessary to adjust the temporal pattern or intensity of optogenetic stimulation.

It remains to be seen whether operant learning can occur by pairing roll or backup behaviour with reward or punishment. Conditioning these actions is challenging given their infrequency in naïve, freely behaving animals. Rolls only occur in response to noxious stimuli (***Ohyama et al., 2013, 2015***; ***Robertson et al., 2013***; ***Tracey et al., 2003***). Backups also occur at very low rates (***Masson et al., 2020***). Consequently, the amount of US which larvae would receive during paired training would be very small, making observable memory formation more difficult. Our high-throughput tracker could potentially address this challenge with probabilistic, thermogenetic activation of roll or back-up command neurons (***Ohyama et al., 2015***; ***Carreira-Rosario et al., 2018***) and optogenetic reward when performing the desired action.

### Neural circuits of operant conditioning

From the available data, it cannot be concluded that the brain and SEZ are dispensable for operant conditioning in *Drosophila* larvae. Examples from both vertebrates (***Grau et al., 1998***) and invertebrates (***Horridge, 1962***; ***Booker and Quinn, 1981***) show the spinal cord or VNC as sufficient for learning, suggesting that conserved mechanisms exist for brain-independent operant conditioning across species. This does not, however, exclude the possibility that there exist alternative learning pathways using the brain. In mammals (***Redgrave et al., 2011***; ***Balleine et al., 2009***) and birds (***Fee and Goldberg, 2011***), brain correlates of operant conditioning have been identified. It is unclear where such pathways would be located in the insect brain. Both our larval experiments and previous adult fly studies (***Colomb and Brembs, 2016***; ***Wolf et al., 1998***; ***Colomb and Brembs, 2010***; ***Booker and Quinn, 1981***) support the idea that operant conditioning can occur independently of the MB, such that other learning centres might exist. To determine whether larval operant conditioning can be fully mediated by the VNC or whether the brain or SEZ are necessary, new driver lines must be created. A collection of sparse split-GAL4 lines, each specific to a distinct group of serotonergic neurons, could help identify the minimum subset of neurons necessary for conveying a US in our bend direction paradigm.

Even if the learning signal for operant conditioning can be mapped to a few serotonergic neurons, there remain several open questions regarding the neuronal mechanisms underlying this learning. Locally, neurons could drive synaptic plasticity or modulate the intrinsic excitability of their postsynaptic partners. Alternatively, the learning signal could propagate further downstream, yielding learning correlates elsewhere in the network. Furthermore, memory formation requires integrating the US with information about the occurrence of the reinforced action. Motor feedback (e. g. efference copy, ***Webb, 2004***; ***Fee, 2014***) or proprioceptive input could be used to transmit this movement signal to higher-level circuits for convergence with the valence-encoding US. However, if memory formation occurred at a lower level, the action-specific signal and associated valence could be locally integrated inside the motor or premotor neuron without the need for feedback loops.

***Lorenzetti et al. (2008)*** proposed intracellular mechanisms for modulating the intrinsic excitability of the *Aplysia* premotor neuron B51 during operant conditioning, mediated by the highly conserved protein kinase C (PKC) gene. PKC signalling is also essential for operant conditioning in *Lymnaea* (***Rosenegger and Lukowiak, 2010***) and adult *Drosophila* (***Brembs and Plendl, 2008***; ***Colomb and Brembs, 2016***). If the *Drosophila* larva showed evidence of PKC-induced motor neuron plasticity, EM reconstruction of the pathways between the serotonergic neurons of the VNC and the PKC-positive motor neurons could further elucidate the mechanisms of memory formation and retrieval. A similar investigation of the *Drosophila* gene *FoxP* may also be informative, as its mutation in the adult results in impaired operant self-learning (***Mendoza et al., 2014***). The vertebrate homologue *FOXP2* is associated with deficits in human speech acquisition (***Lai et al., 2001***), song learning in birds (***Haesler et al., 2007***), and motor learning in mice (***Groszer et al., 2008***).

### Serotonin as a learning signal

A limited set of studies have shown that serotonergic signalling is sufficient to induce learning (***Liu et al., 2014***; ***Hawkins and Byrne, 2015***; ***Ganguly et al., 2020***). Previous *Drosophila* studies highlight other roles of serotonin in associative learning (***Yu et al., 2005***; ***Keene et al., 2004, 2006***; ***Lee et al., 2011***; ***Huser et al., 2017***). ***Sitaraman et al.*** (***2012***) have shown that synaptic transmission from serotonergic neurons is essential for appetitive olfactory conditioning in the adult. Aversive olfactory memory formation is impaired in flies fed with a tryptophan hydroxylase inhibitor which blocks serotonin biosynthesis (***Lee et al., 2011***). Furthermore, serotonin receptor signalling is required for memory formation in classical conditioning tasks (***Johnson et al., 2011***). In larvae, aversive olfactory conditioning is impaired by either ablation of serotonergic neurons during development or mutations in a serotonin receptor gene (***Huser et al., 2017***).

Our work suggests a novel role of serotonin as a reward signal for learning in *Drosophila* larvae. In our olfactory classical conditioning screen, optogenetic stimulation of serotonergic neurons in the brain and SEZ was sufficient to induce strong appetitive learning. Conversely, operant conditioning necessitated serotonergic neuron activity in the VNC. Since it remains unclear to what extent serotonergic neurons in the brain and SEZ are also involved in the operant conditioning effect we observed, it is possible that some neurons mediate both forms of associative learning.

Further investigation is necessary to better understand the function of serotonin in memory formation. It is possible that even a single instance of learning leads to a variety of changes across the nervous system. In the case of operant conditioning, higher brain centres, motor command neurons, premotor circuits and motor neurons would all qualify as potential learning sites. In addition to thoroughly analysing the expression patterns of driver lines used in our classical conditioning screen, developing new, sparse driver lines targeting serotonergic neurons would be valuable for identifying the minimal subset of neurons which provide the serotonergic learning signal. The larval connectome could be used to subsequently trace the paths from these neurons to the MB. One could then test whether learning as induced by the serotonergic US remains intact when these connections are silenced. The expression pattern of serotonin receptors could also provide clues about how the serotonergic signal triggers learning. One should certainly consider the possibility that learning is not induced by serotonin itself, but by other neurotransmitters coexpressed by serotonergic neurons. This could be assessed by suppressing serotonin biosynthesis in the desired neuronal subset.

## Materials and methods

### High-throughput closed-loop tracker

#### Hardware set-up

A high-resolution camera (3072 × 3200 pixels) (#TEL-G3-CM10-M5105, Teledyne DALSA, Ontario, Canada) positioned above a 23 cm × 23 cm 4% agarose plate captured 8-bit greyscale images at 20 Hz. The agarose plate was illuminated from below by a 30 cm × 30 cm 850 nm LED backlight (#SOBL-300×300-850, Smart Vision Lights, Norton Shores, Michigan) equipped with intensity control (#IVP-C1, Smart Vision Lights, Norton Shores, Michigan). An 800 nm longpass filter (#LP800-40.5, Midwest Optical Systems, Palatine, Illinois) mounted on the camera blocked all visible wavelengths, including those used for optogenetics. When the agarose plate comprised most of the camera image, each pixel corresponded to a 72.92 µm diameter section of the plate.

Each camera image was processed in parallel on both the host computer (#T7920, running Windows 10, Dell Technologies Inc, Round Rock, Texas) and an FPGA device (#PCIe-1473R-LX110, National Instruments, Austin, Texas). LabVIEW 2017 (National Instruments, Austin, Texas) software extracted larval contours and interfaced with C++ software that performed real-time behaviour detection. The LabVIEW software controlled closed-loop optogenetic and thermogenetic stimulation in response to these detected behaviours.

#### Multi-animal detection and tracking

Raw camera images were read by the FPGA at 20 Hz and then sent to the host computer. The Lab-VIEW process on the host computer then filtered out non-larval objects by combining background subtraction and binary thresholding. The remaining objects were each enclosed in a rectangular box of minimal size, with edges parallel to the camera image axes. We defined the following criteria to detect third-instar larvae within these boxes:

- Pixel intensity range (default 25–170): the minimum and maximum brightness values for pixels selected by binary thresholding (between 0 and 255 for an 8-bit image).
- Box side length (pixels) (default 6–100): the range of eligible values for width and height of each box.
- Box width + height (pixels) (default 12–200): the range of eligible values for the sum of each box’s width and height.
- Box area (pixels) (default 300–900): the range of eligible values for the area of each box.

To track larvae over time, the host computer assigned a numerical identifier to each eligible object. We used distance-based tracking with a hard threshold of 40 pixels to maintain larval ID based on centroid position. Although identity was lost when larvae touched or reached the plate’s edge, new IDs were generated when larvae matched detection criteria. For each of the largest 16 objects, the host computer sent a binary pixel pattern and location (defined as the centre of the box) to the FPGA. Since the host computer required more than 50 ms of run time for object detection, this process was not executed in every frame. On average, the FPGA received updated objects and their locations every three frames.

The FPGA extracted object contours in three steps. Within a 2 cm2 region of interest around the object’s centre, the FPGA first applied a user-defined binary threshold, then applied both vertical and a horizontal convolution with a 2 x 1 XOR kernel, and finally generated edge pixels by combining the results of the two convolutions using an OR operation. Contours were extracted from edge images using the Moore boundary tracing algorithm (***Gonzalez and Woods, 2018***) with three added error capture procedures. First, if the algorithm yielded a contour that ended prematurely or contained small loops, the construction process could be reversed by up to 16 contour points to find an alternative contour. Second, 10,000 FPGA clock cycles (≈ 100 us) was the maximum allotted execution time, with each pixel comparison occurring within one clock cycle. In the rare event that this window was exceeded, the algorithm returned the already constructed contour points. Third, a contour containing fewer than 63 points was rejected and the FPGA returned the last valid contour detected for a given larva ID. The algorithm stopped when none of the remaining neighbours were edge pixels (***Figure 1***–***Figure Supplement 1***).

#### Contour processing and landmark detection

An undesired result of the FPGA contouring algorithm was the variable number of contour points across larvae and frames. We aimed to detect behaviour based on a smooth contour with a fixed number of 100 contour points. This contour regularization was achieved inside the Behaviour Programme using Fourier decomposition and reconstruction as in ***Masson et al. (2020***).

The initial detection of head and tail was implemented on FPGA. The larva’s head and tail were defined as the contour points with the sharpest and second-sharpest curvature, respectively (***Figure 1***–***Figure Supplement 2***). While correct in most cases, this calculation sometimes led to flipped detection of the two body ends. The Behaviour Programme flagged and corrected these false detection events at run time by calculating the distance head and tail traveled between frames and tracking the number of correct versus flipped detection events. The vote system correction commonly failed when the larva made large angle bends. The resulting contour was nearly-circular and exhibited similar curvature across all points. The solution required resetting the vote tallies when detecting these ball events (***Figure 1***–***Figure Supplement 2***).

We defined the larval spine as 11 points running along the central body axis from `head to tail` (***Figure 1***–***Figure Supplement 3***; ***Swierczek et al., 2011***). In addition to `head` and `tail`, the Behaviour Programme calculated three equally distributed landmark points along the spine (`neck_top, neck`, and `neck_down`). A fourth landmark, the `centroid`, defined the larva’s location. The six landmarks were collectively used to extract features for training behaviour classifiers (***Figure 1***–***Figure Supplement 3***).

The Behaviour Programme transformed the raw contour and spine from camera coordinates (in pixels) to world coordinates (in mm). If stable larval detection criteria were met, all spine points were temporally smoothed using exponential smoothing (***Figure 1***–***Figure Supplement 3***).

#### Feature extraction

We developed a machine learning approach to address the high deformability of the larva shape, ensure live execution, reduce overfitting, and limit the volume of data tagging. What follows is a brief summary of larval features describing motion direction, body shape, and velocity that were calculated from the contour and spine data inside the Behaviour Programme. Features were designed as in ***Masson et al. (2020)***, with notable modifications required to run the inference live:

1. Motion Direction (***Figure 1***–***Figure Supplement 4)***

- `direction_vector`: normalised vector describing the main body axis
- `direction_head_vector`: normalised vector describing the head axis
- `direction_tail_vector`: normalised vector describing the tail axis
2. Body Shape (***Figure 1***–***Figure Supplement 5)***

- `skeleton_length`: summed distances between consecutive spine points
- `perimeter`: summed distances between neighbouring contour points
- `larva_arc_ratio`: ratio of contour perimeter to convex hull perimeter (`larva_arc_ratio` ≥ 1 and was close to 1 when larva was in either straight or ball-like shape)
- `larva_area_ratio`: ratio of the areas enclosed by the contour and its convex hull (0 ≤ `larva_area_ratio` ≤ 1 and was close to 1 when the larva was in either straight, heavily curved, or ball-like shape)
- `eig_reduced`: 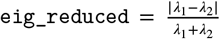 where *λ*_1_, *λ*_2_ were the eigenvalues of the structure tensor of the larval contour with respect to the `neck`(0 ≤ `eig_reduced` ≤ 1 and `eig_-reduced` decreased as the bend amplitude of the larva increased)
- `s`: normalised angle along the body (−0.5 ≤ `s` ≤ 1, was close to 1 when larva was straight, and decreased with increasing bend amplitude)
- `asymmetry`: sine of the angle between `direction_vector` and `direction_head_vector` (`asymmetry` > 0 when larva bent left and `asymmetry` < 0 when larva bent right)
- `angle_upper_lower`: absolute angle between `direction_vector` and `direction_head_-vector` (despite similarity to `asymmetry`, this develops different dynamics following temporal smoothing, which are valuable for stable left and right bend detection)
3. Velocity (***Figure 1***–***Figure Supplement 6)***

- Velocity of all six landmark points (`head_speed, neck_top_speed, neck_speed, neck_down_-speed, tail_speed`, and `v_centroid`) in mm/s over interval *dt* = 0.2 *s* (four frames)
- `v_norm`: arithmetic mean of `neck_top_speed, neck_speed`, and `neck_down_speed`, passed through a hyperbolic tangent activation function to suppress excessively large values
- `speed_reduced`: relative contribution of `neck_top_speed` to `v_norm`, passed through a hyperbolic tangent activation function to suppress excessively large values (`speed_reduced` increased when the anterior larval body moved quickly compared to the posterior, e. g. when a bend was initiated)
- `damped_distance`: distance (mm) travelled by `neck`, giving greater weight to recent over past events
- `crab_speed`: lateral velocity (mm/s), defined as the component of `neck_speed` orthogonal to `direction_vector_filtered`
- `parallel_speed`: forward velocity (mm/s), defined as the component of `neck_speed_-filtered` parallel to `direction_vector_filtered`
- `parallel_speed_tail_raw`: tail’s forward velocity (mm/s), defined as the component of `tail_speed_filtered` parallel to `direction_tail_vector_filtered`
- `parallel_speed_tail`: similar to `parallel_speed_tail_raw`, with the difference that `tail_-speed_filtered` was normalised prior to calculating the dot product (i. e. a measure of tail movement direction which took values between −1 (backward) and +1 (forward))

To extract features in real time and address various sources of noise, we implemented exponential smoothing defined as follows for a given feature `f` (***Figure 1***–***Figure Supplement 7)***:

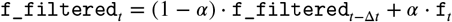

where t is unitless, but derived from the experiment time in seconds, 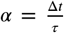 with Δ*t* = 0.05 *s* and *τ* = 0.25 *s*. Features that had the potential to exhibit large value deviations (e. g. `v_norm`) were instead bounded using a hyperbolic tangent function. Additionally, some features were exponentially smoothed over a longer time window (where 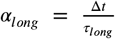 with Δ*t* = 0.05 *s* and *τ*_*long*_ = 5 *s*) (*Figure 1*–*Figure Supplement 7*).

Convolution was used to approximate a smoothed squared derivative for each feature (***Figure 1***–***Figure Supplement 8)***; useful for integrating information over time without needing to further expand the feature space. The underlying mathematical concepts were motivated by ***Masson et al. (2012)***. For a given feature `f` at time *t*, f_convolved_squared was calculated as follows:

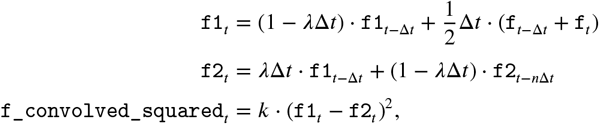

where Δ*t* = 0.05 *s*, 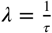, *τ* = 0.25*s* and *n* = 5 *s*. *k* values were empirically chosen for each feature.

#### Behaviour classifiers

Behaviour classifiers were developed using a user interface similar to JAABA (***Kabra et al., 2013)***. The underlying algorithms combined trained neural networks and empirically determined linear thresholds. We developed a MATLAB (MathWorks, Natick, Massachusetts) user interface with functions for data visualisation, manual annotation, and machine learning using the Neural Network Toolbox, the Deep Learning Toolbox, and the Statistics and Machine Learning Toolbox. Here we briefly describe the behaviour classifiers and provide performance results based on manual validation (***Table 1)***.

**Table 1.**
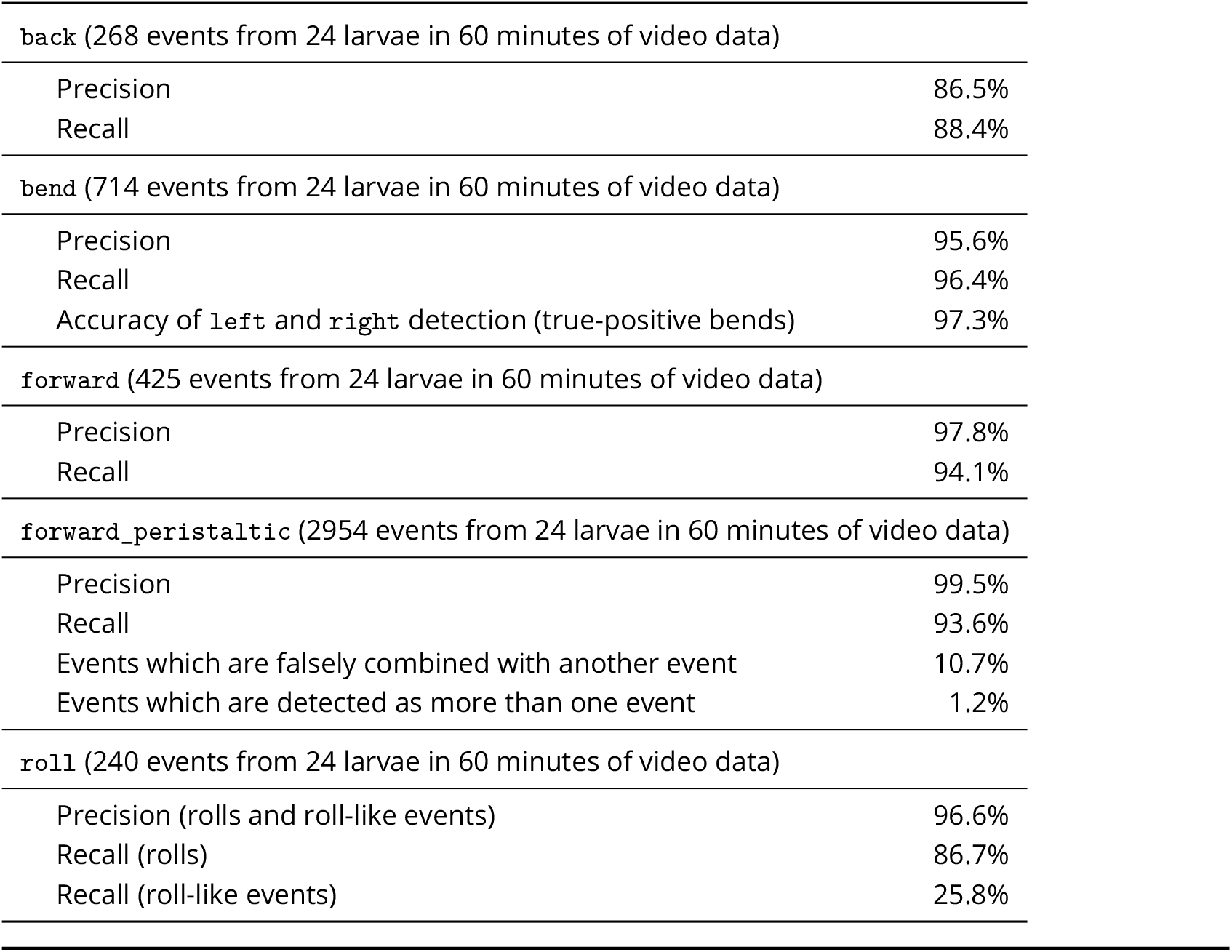
Manual quantification of behaviour detection performance.

The `bend` classifier was based on predefined thresholds for temporally smoothed body shape features and was itself exponentially smoothed over time. Independent `left` and `right` classifiers were used to initially detect bend direction. To detect left and right bends, these classifiers were combined with the smoothed bend classifier using an AND conjunction. The raw time series of left and right bends was further smoothed post-acquisition using a custom MATLAB script: two bends to the same side separated by less than 200 ms were combined into a single long bend, and short bends of less than 200 ms were removed from analysis.

To improve `left` and `right` detection performance, we developed a classifier for circular larval contours. This ball classifier used a feed-forward neural network with a single fully connected hidden layer whose inputs were normalised values of `eig_reduced, larva_arc_ratio, and larva_-area_ratio`. The hidden layer consisted of five neurons with a hyperbolic tangent activation function. The output layer contained a single neuron and used a sigmoid activation function. The neural network was trained in MATLAB on a manually annotated data set for 500 epochs using a cross-entropy loss function and scaled conjugate gradient backpropagation. If a `ball` was detected within the previous 1.5 s, `left` and `right` classifiers were overwritten to match the last detected bend direction prior to the beginning of the `ball`.

The `back` classifier detected individual backward peristaltic waves based on thresholds for smoothed tail velocity features combined with no `ball` detection within the previous 1.5 s.

Two different classifiers were used to detect crawling. `forward` detected longer forward crawl periods based on thresholds for smoothed tail velocity features combined with no `ball` detection within the previous 1.5 s. `forward_peristaltic` detected individual forward peristaltic waves based on the `forward` classifier and a threshold on forward tail velocity.

The `roll` classifier was based on thresholds for body shape and velocity combined with no `ball` detection and was exponentially smoothed over time. If a `roll` was detected within the previous 1.5 s, `forward, forward_peristaltic`, and `back` classifier values were reset to reduce false-positive detection for these classifiers. Unusual behaviour patterns such as rapid bending or twitching could be observed in addition to true larval rolling. These behaviours were considered “roll-like” events during manual validation of the roll classifier’s performance.

#### Optogenetic stimulation

Optogenetic stimulation was achieved using two digital micromirror devices DMDs to project light patterns onto larvae on the agarose plate. During the hardware design process, two different DMD models were tested. One contained an integrated 613 nm LED (#CEL-5500-LED, Digital Light Innovations, Austin, Texas) and the other (#CEL-5500-FIBER, Digital Light Innovations, Austin, Texas) received input from an external 625 nm LED (#BLS-GCS-0625-38-A0710, Mightex Systems, Ontario, Canada) controlled by a BioLED light source control module (#BLS-13000-1, Mightex Systems, Ontario, Canada) and fed through an optic fibre (#LLG-05-59-420-2000-1, Mightex Systems, Ontario, Canada). Both DMDs operated like a 768 × 1024 pixel monochrome red light projector with numerous rotatable micromirrors used to modulate the intensity of individual pixels. Although both achieved similar light intensities, each DMD on its own was insufficient for optogenetic stimulation of larvae. We installed both devices on the system such that their projections each covered the entire agarose plate. In this way, the summed light intensities of the two DMDs could be achieved at all locations. Accurately aiming light at crawling larvae required spatial calibration of each DMD. Calibration was performed by projecting square spots at fixed DMD pixel locations and linearly fitting the corresponding camera coordinates.

We determined that DMD illumination using the default light output was not uniform at plate level, which could have resulted in variable optogenetic stimulation depending on larval location. The maximum achievable light intensity at the plate’s edge was approximately 40% of the peak value at its centre. We therefore normalised the pixel intensity of the DMD image to the highest intensity uniformly achievable at all plate locations. A look-up table containing the normalisation factor for each DMD pixel was then calculated using bi-linear interpolation with approximately 100 light intensity values measured across the plate. To accommodate for possible differences in non-uniformity between the two DMDs, this intensity calibration was performed for both DMDs simultaneously following spatial calibration. When fully calibrated, the system could achieve a uniform light intensity of 285 μW/cm^2^.

A user-defined Behaviour Programme protocol operated on the behaviour detection output and sent 8-bit optogenetic stimulation instructions to the LabVIEW application. Because the Lab-VIEW application updated DMD projections at 20 Hz, the delay between behaviour detection and closed-loop optogenetic stimulation of individual larvae did not exceed 50 ms. Furthermore, if two or more larvae were close enough such that their corresponding stimulation areas overlapped, the light intensity in the overlapping region was set to the smallest of those values to avoid undesired stimulation.

#### Thermogenetic stimulation

Thermogenetic stimulation was achieved by heating up larvae with a custom IR laser set-up. A 1490 nm laser diode beam (#2CM-101, SemiNex, Peabody, Massachusetts) was fed into a two-axis galvanometer system (#GVSM002, Thorlabs, Newton, New Jersey), both controlled by an analogue output device (#PCIe-6738, National Instruments, Austin, Texas). Two mirrors inside the galvanometer were rotated around orthogonal axes to target the beam spot to any user-defined location on the agarose plate. The beam spot measured approximately 5 mm in diameter, depending on the beam’s angle of incidence to the plate. Mirror positions were controlled by two integrated motors that received voltage inputs. Each voltage pair clearly defined the laser beam’s position.

Spatially calibrating the galvanometer was necessary to obtain a map between larval locations in world coordinates and the mirror motor input voltages. A visible aiming beam was scanned across the agarose plate using a fixed set of voltage pair inputs to the galvanometer. With the optical filter removed from the camera, the aiming beam’s location in camera coordinates was automatically extracted from the image using binary thresholding. Two voltage-to-camera look-up tables were generated through bi-linear interpolation of these measured coordinates. For accurately targeted thermogenetic stimulation, the location of the larval centroid was first converted to camera coordinates using the existing world-to-camera transform and was then mapped to a pair of galvanometer input voltages using the look-up tables.

Laser intensity calibration was also necessary to ensure that all larvae received the same stimulation regardless of their position on the agarose plate. A larva’s location changed the laser beam’s angle of incidence, causing the illuminated spot at plate level to take an elliptical shape with variable size. Although laser beam power was constant, the changing spot area generated inconsistencies in the amount of IR light covering each larva. Calibration was used to normalise the desired laser intensity to achieve constant power per unit area. A visible aiming beam was scanned across the plate and the camera image automatically measured the beam’s spot size at various locations. Bi-linear interpolation was then used to generate a pixel-wise look-up table containing the necessary scaling factors for the laser power. At the location where the laser spot area was smallest, the maximum power was reduced to 67.3%. We also accounted for a nonlinear relationship between the laser source input voltage and the laser’s total power output by generating a voltage-to-power map from manual measurements. With these transformations, the system could calculate the laser source input voltage necessary to produce uniform, 5.26 W stimulation at any location.

A user-defined Behaviour Programme protocol operated on the 20 Hz behaviour detection output and sent thermogenetic stimulation instructions to the LabVIEW application which controlled the galvanometer and laser. Four centroid locations were specified on every frame, enabling a single galvanometer to cycle the laser beam between four individual larvae at 20 Hz. Within the available 50 ms time window, each larva was heated for 11 ms. Switching off the laser input for 1.5 ms between larvae accounted for small time fluctuations surrounding each new galvanometer position update and helped avoid undesired stimulation of other plate areas (***Figure 2D)***. If fewer than four objects were detected in a given frame, the remaining galvanometer target locations were set to the plate’s centre and the corresponding laser intensity was set to zero. This temporal pattern of galvanometer position updates yielded no more than 100 ms delay between behaviour detection and closed-loop thermogenetic stimulation.

Three parameters influenced larval temperature increase following thermogenetic stimulation with the IR beam: i) the laser power, ii) the total duration of the stimulus, and iii) the order in which the galvanometer cycles between locations in its 80 Hz movement. Preliminary experiments suggested that these parameters could be adjusted to simultaneously stimulate eight or twelve larvae using a single galvanometer. This could potentially eliminate the need to install three additional laser sources to target all 16 larvae.

### Software availability

All software code is available upon request.

### Fly strains and larval rearing

We used the following fly strains: *58E02-Gal4* (Bloomington stock 41347), *69F06-Gal4* (Bloomington stock 39497), *72F11-Gal4* (Bloomington stock 39786), *attP2* (***Pfeiffer et al., 2008)***, *Ddc-Gal4* (***Li et al., 2000)***, *SS01989* (own stock), *TH-Gal4* (***Friggi-Grelin et al., 2003)***, *Tph-Gal4* (***Park et al., 2006)***, *Trh-Gal4* (***Alekseyenko et al., 2010)***, *UAS-CsChrimson* (Bloomington stock 55134), *UAS-CsChrimson; tsh-LexA, LexAop-Gal80* (Dr Stefan Pulver, Dr Yoshinori Aso), *UAS-dTrpA1* (Dr Paul Garrity), *UAS-GFP* (***Nern et al., 2015)***, and *w1118* (***Hazelrigg et al., 1984)***.

Fly stocks were maintained in vials filled with standard cornmeal food (***Wirtz and Semey, 1982***; 49.2 ml of molasses, 19.9 g of yeast, 82.2 g of cornmeal, 7.4 g of agarose, 9.8 ml of 20% Tegosept solution in 95% ethanol and 5.2 ml of propionic acid in 1 litre of water). For proof-of-principle and operant and classical learning experiments, eggs were collected overnight for approximately 12–18 hours on standard cornmeal food plates with additional dry yeast to increase laying. These experiments were performed using foraging-stage third-instar larvae (72–96 hours after egg laying) reared at 25°C and 65% humidity (***Ohyama et al., 2013, 2015***; ***Jovanic et al., 2016, 2019***; ***Eschbach et al., 2020b)***. Specifically for optogenetics experiments, larvae were raised in the dark and a 1:200 retinal solution (diluting 1 g of powdered all-*trans*-retinal (#R240000, Toronto Research Chemicals, Ontario, Canada) in 35.2 ml of 95% ethanol) was added to the food unless indicated otherwise. For immunohistochemistry, eggs were collected during daytime for approximately four hours on standard cornmeal food plates with added yeast. Dissections were performed using wandering-stage third-instar larvae (118–122 hours after egg laying).

### Immunohistochemistry and confocal imaging

All dissections, immunohistochemical stainings, and confocal imaging were done using a procedure adapted from ***Jenett et al. (2012)*** and ***Li et al. (2014)***. Larval CNSs were dissected in cold 1x phosphate buffer saline (PBS, Corning Cellgro, #21-040) and transferred to tubes filled with cold 4% paraformaldehyde (Electron Microscopy Sciences, #15713-S) in 1x PBS. Tubes were incubated for one hour at room temperature. The tissue was then washed four times in 1x PBS with 1% Triton X-100 (#X100, Sigma Aldrich St. Louis, Missouri) (PBT) and incubated in 1:20 donkey serum (#017-000-121, Jackson Immuno Research, West Grove, Pennsylvania) in PBT for two hours at room temperature.

The tissue was then incubated in the primary antibody solution, first for four hours at room temperature and then for two nights at 4°C. This solution contained mouse anti-Neuroglian (1:50, #BP104 anti-Neuroglian, Developmental Studies Hybridoma Bank, Iowa City, Iowa), rabbit anti-green fluorescent protein (GFP) (1:500, #A11122, Life Technologies, Waltham, Massachusetts) and rat anti-N-Cadherin (1:50, #DN-Ex #8, Developmental Studies Hybridoma Bank, Iowa City, Iowa) in PBT. This solution was then removed and the tissue washed four times in PBT. The tissue was then incubated in the secondary antibody solution, first for four hours at room temperature and then for two nights at 4°C. This solution contained Alexa Fluor 568 donkey anti-mouse (1:500, #A10037, Invitrogen, Waltham, Massachusetts), FITC donkey anti-rabbit (1:500, #711-095-152, Jackson Immuno Research West Grove, Pennsylvania) and Alexa Fluor 647 donkey anti-rat (1:500, #712-605-153, Jackson Immuno Research West Grove, Pennsylvania) in PBT. After removal of the secondary solution, the tissue was washed in PBT four times and mounted on a coverslip coated with poly-L-lysine (#P1524-25MG, Sigma Aldrich, St. Louis, Missouri).

The coverslip with the CNSs was dehydrated by moving it through a series of jars containing ethanol at increasing concentrations (30%, 50%, 75%, 95%, 100%, 100%, 100%) for ten minutes each. The tissue was then cleared by soaking the coverslip with xylene (#X5-500, Fisher Scientific, Waltham, Massachusetts) three times for five minutes each. Finally, the coverslips were mounted in dibutyl phthalate in xylene (DPX, #13512, Electron Microscopy Sciences, Hatfield, Pennsylvania) with the tissue facing down on a microscope slide with spacers. The DPX was allowed to dry for at least two nights prior to confocal imaging with an LSM 710 microscope (Zeiss).

Details on the confocal imaging settings are provided in the respective figure captions. Confocal images were analysed using Fiji (ImageJ). Neurons were counted by specifying regions of interest around the cell bodies using raw image stacks.

### Verification of optogenetic and thermogenetic stimulation eficiency

We assessed the multi-larva tracker’s optogenetic and thermogenetic stimulation effciency through open-loop experiments. The behavioural readout was rolling upon exposure to stimulation. All larval handling and experiments were performed in the dark to avoid unintended optogenetic stimulation. The one-minute experiment protocol began with a 15 s initialisation period in which larvae acclimated to the agarose plate and the `roll` behaviour classifier stabilised. In three subsequent 15 s stimulation cycles, larvae received 5 s of open-loop stimulation followed by 10 s without stimulation (***Figure 2B, Figure 2E)***. Optogenetics were performed with the maximum available red light intensity of 285 μW/cm^2^. Thermogenetics were performed with 40% of the maximum available laser intensity.

We analysed both optogenetic and thermogenetic experiment data using identical assessment and exclusion criteria. For each larva, the criterion for a single `roll` was detection of the behaviour for at least 300 ms during a given 15 s stimulation cycle. This threshold ensured true rolls were counted, as opposed to rapid larval bends characteristic of aversion to light.

### High-throughput operant conditioning

#### Experiment procedures

We performed high-throughput operant conditioning using our multi-larva closed-loop tracker. All larval handling and experiments were performed in the dark to avoid unintended optogenetic stimulation. We used water to wash approximately 10–12 larvae out of their food. Using a brush, we immediately placed these larvae into the centre of the agarose plate in such a way that they were not touching each other. We placed the agarose plate inside the tracker on top of the backlight and then shut the tracker door. Larvae were given at least 30 s to accustom to their new environment before we started the experiment.

The experiment protocol began and ended with a one-minute test period without optogenetic stimulation. Between these test periods were four, three-minute training sessions during which larvae received red light stimulation of 285 μW/cm2 for the entire duration of the detected bend. Which side received stimulation was randomized across trials such that approximately 50% of larvae were trained to develop a right bend preference and 50% a left bend preference. No stimulus was triggered when the larva was bending right or when its body was straight. The test periods were each separated by three-minute periods without stimulation. After the first minute of this period, a brush was used to gently move all larvae back to the centre of the plate and larvae were given time to recover before the beginning of the next training session. This recentring addresses problems encountered when performing prolonged experiments with freely behaving larvae on a small agarose plate. The longer larvae are left undisturbed, the more likely they are to touch the plate’s edge, causing tracking disruption and temporary loss of valid objects. This shrinks sample size and reduces training efficiency by decreasing the proportion of animals which are receiving the stimulus.

Control experiments were designed so that valid objects received optogenetic stimulation uncorrelated with behaviour. These control experiments were split into 60 s time bins, during which each valid object was randomly assigned a stimulus train from this same time bin, pulled from a prior experiment where stimulation correlated with behaviour.

#### Data analysis

Data analysis was conducted using custom MATLAB software. To ensure high quality data, it was necessary to remove invalid objects from the data set prior to behavioural analysis. These included corrupted objects (e.g. scratches on the plate or residual food) that the software mistook for larvae. They also included larvae that lost their object identity and were consequently detected for only part of the experiment (e.g. after temporarily reaching the plate’s edge or touching other larvae). After equally splitting each experiment into 60 s time bins, we retained objects for analysis that fulfilled strict criteria: i) the object must have been detected in every frame of the bin; ii) the object’s initial detection must have occurred at least 20 s prior to the start of the bin; iii) at no point during the bin did the smoothed velocity of the larval centroid exceed 1.5 mm/s; and iv) the mean of the smoothed centroid velocity across the object’s detection period in the bin was at least 0.5 mm/s. To quantify the accuracy of this method, we manually reviewed 350 videos of objects flagged as valid for a given 60 s bin. In this group, we observed no severely corrupted objects. In one case (0.3%), a larva briefly touched another larva. In another case (0.3%), head and tail of a larva were falsely detected the majority of the time, leading to flipped detection of left and right bends.

When analysing valid bin data for operant conditioning of bend direction preference, we counted, for each larva, the numbers of left and right bends initiated within the bin. This was defined as the bend rate towards the respective direction. The probability of the larva bending towards the side paired with the optogenetic stimulus was defined as the ratio of the number of bends towards this side to the total number of bends initiated in the bin. We pooled together all larval data within each bin because bends to the left and right were each paired with the optogenetic stimulus for approximately half of the larvae. Mean and standard error were calculated for bend rate to the stimulated side, bend rate to the unstimulated side, and probability of bending towards the stimulated side. For the control condition in which larvae received random stimulation during 50% of bends regardless of direction, we calculated mean and standard error for bend rates to the left and right and the probability for bending towards the left. Bend rates to either side were compared to each other using a two-sided Wilcoxon signed-rank test. The probability for bending to a given side was compared to chance level (0.5) using a two-sided Wilcoxon signed-rank test. The behaviour characteristics of experimental animals were compared to the control group using a two-sided Mann-Whitney *U* test.

### Classical conditioning

#### Experiment procedures

*CsChrimson* (***Klapoetke et al., 2014)*** was expressed under the control of driver lines targeting candidate valence-conveying neurons. These driver lines were classified based on expression pattern and previous functional data and are known to drive expression in larvae. Optogenetic activation of neurons (US) was paired with odour presentation (CS) to induce olfactory memory (***Figure 4A)***. For each driver line, data was acquired from at least two separate crosses.

Classical conditioning followed a procedure similar to those described in ***Gerber and Hendel (2006)***, ***Saumweber et al. (2011)*** and ***Eschbach et al. (2020b)***. Approximately 40 third-instar larvae were transferred onto a 4% agarose petri dish. Larvae were presented with an odour (1:10^4^ ethyl acetate in ddH_2_O) pipetted onto two pieces of filter paper attached to the lid of the dish. This enclosed dish was exposed to red light (630 nm, 350 μW/cm^2^) for three minutes. Larvae were then transferred to a new agarose-filled petri dish with no odour on its lid (“air”) and placed in the dark for three minutes. This training procedure was repeated three times, with alternating presentation of odour/light and air/dark (paired group). An unpaired group receiving reciprocal stimulus presentation (odour/dark, air/light) was trained simultaneously. This ensured that any observed effects were attributable to learning rather than innate odour preference or avoidance. The training trial order was reversed in half of the experiments, starting with air instead of odour presentation.

After training, larvae of both paired and unpaired groups were immediately transferred to a 1 cm middle zone in the centre of fresh agarose-filled petri dishes. A lid was placed on each dish, with odour presented on one side (odour side) but not the other (air side). After a three-minute test period in the dark, the number of larvae on the odour side, the air side, and in the middle zone were manually counted and entered into an Excel spreadsheet (Microsoft Corporation, Remond, Washington).

#### Data analysis

All data was manually entered into MATLAB and analysed using custom software. For each experiment, a performance index (PI) was calculated as follows:

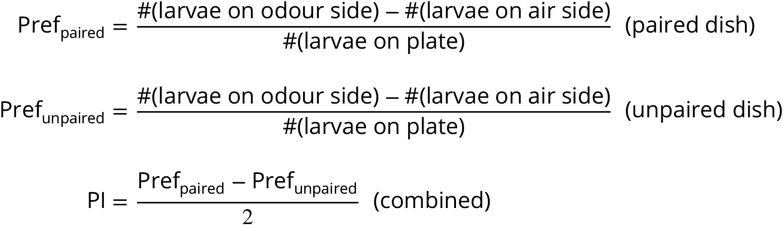

PIs take values between −1 and +1, where a positive PI reflects appetitive learning and a negative PI reflects aversive learning. Mean and standard error were calculated for each condition. Statistical differences between two groups were tested using a two-sided Mann-Whitney *U* test with Bonferroni correction. Significance compared to zero was tested with a two-sided Wilcoxon signed-rank test with Bonferroni correction.

## Acknowledgments

We thank Dr. Chris McRaven for design and technical assistance with the thermogenetic laser light source; Howard Hughes Medical Institute (HHMI) Janelia FlyCore and FlyLight teams for assistance with fly crosses, fly food, and confocal imaging; Gates Cambridge Trust, Cambridge Trust, HHMI Janelia Visiting Scientist Program, University of Cambridge Trinity College, HHMI Janelia, European Research Council, Wellcome Trust, and Medical Research Council for funding.

---

**Figure 1–Figure supplement 1.**
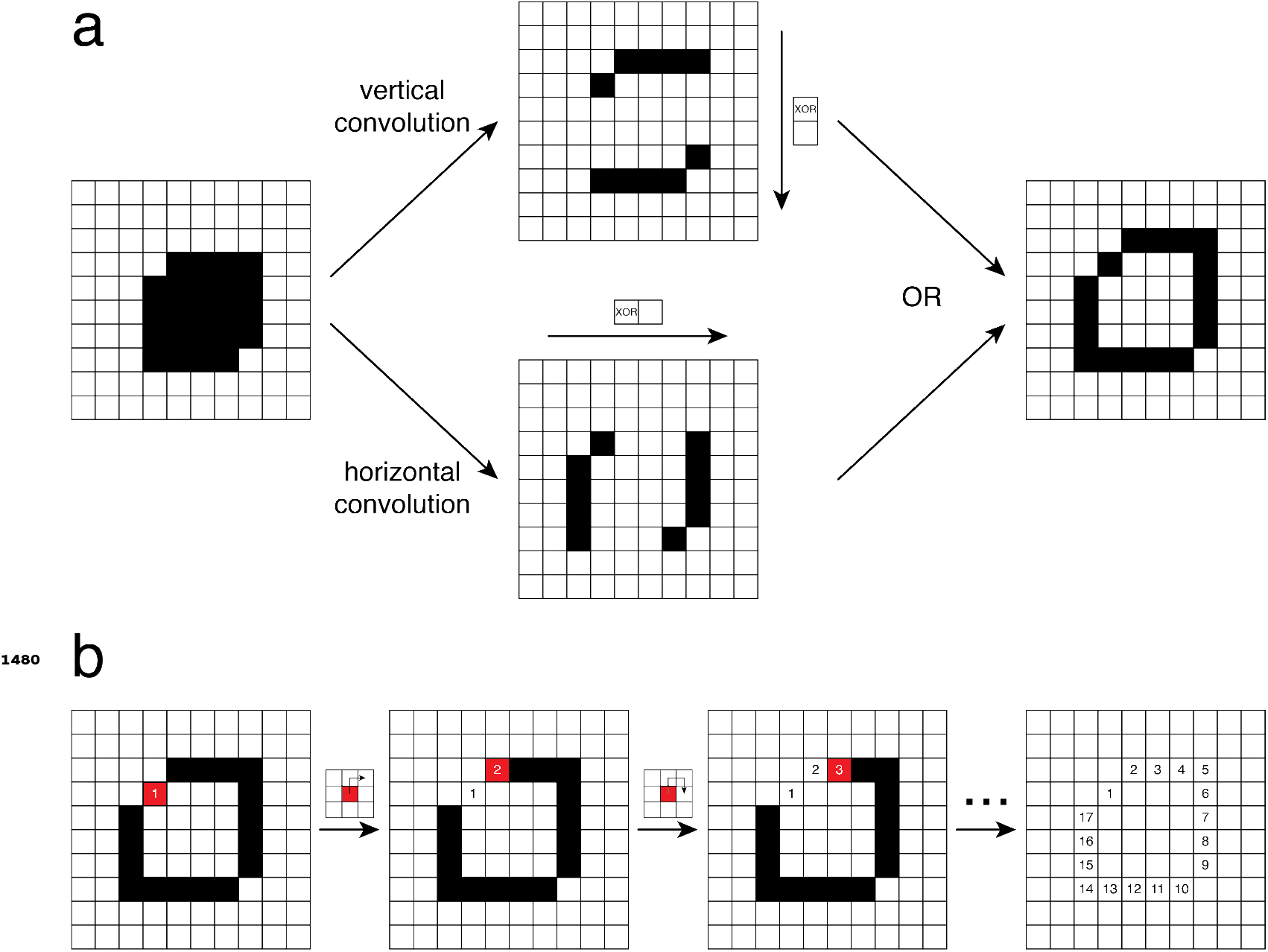
Contour calculation on field-programmable gate array (FPGA). A simplified example is shown using a 10 × 10 pixel box containing a small object. **a.** The object (black) was detected against the background (white) using binary thresholding. Edge pixels were detected by combining the results of vertical and horizontal image convolution with a 2 × 1 XOR kernel using an OR operator. **b.** The contour points were reconstructed in an iterative process, starting with the edge pixel closest to the centre of thebox. The next contour point was defined as the first neighbouring pixel that was found to be an edge pixel. Neighbouring pixels were assessed clockwise from the pixel directly above the contour point. The process ended when no eligible edge pixels could befound.

**Figure 1–Figure supplement 2.**
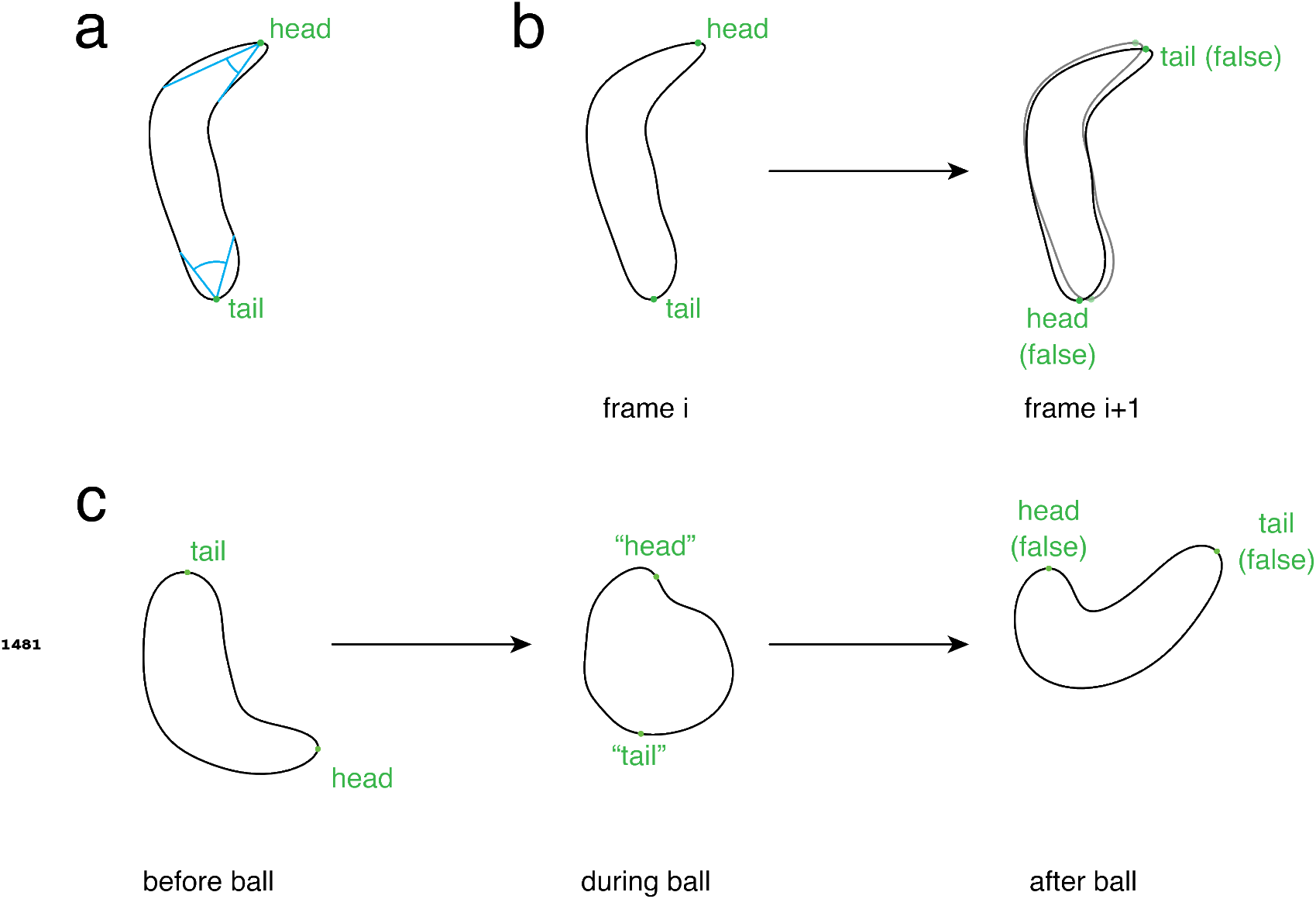
Detecting head and tail. The larval contour (black outline) and head and tail (green) are shown. **a.** Initial detection of head and tail. The head was the contour point with the sharpest curvature. The tail was the contour point with the next-sharpest curvature which did not lie in close proximity to the head. **b.** The initial detection of head and tail was incorrect in some cases. False detection could be corrected by swapping head and tail, thereby minimising the distances from head and tail in the current frame (solid contour) to head and tail in the previous frame (transparent contour). **c.** The correction described in **b** failed if larvae curled up such that the contour appeared circular (“ball”). To eliminate this source of false head and tail detection, these events were detected using a `ball` classifier.

**Figure 1–Figure supplement 3.**
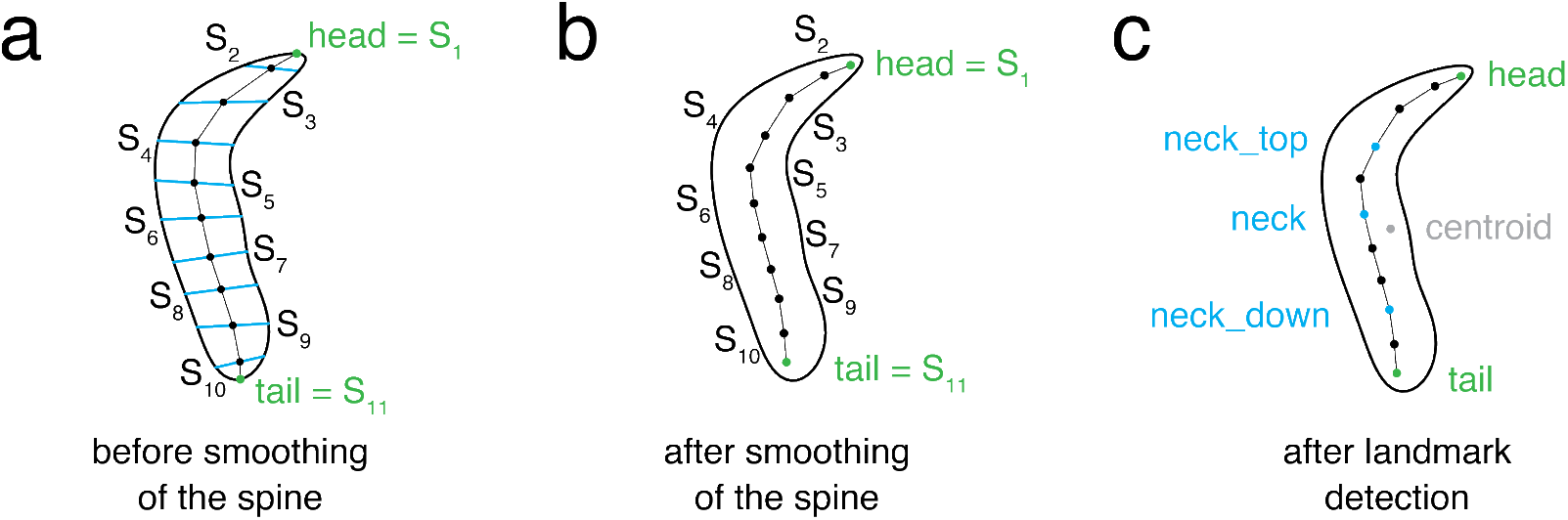
Calculating a smooth spine and landmark points. The larval contour is shown (black outline). The spine ***S*** was comprised of eleven points (black), including head and tail (green). **a.** The raw spine points were obtained by finding the centres between equally spaced contour points on either half of the contour as defined by head and tail. The first spine point was the head, the last spine point was the tail. **b.** The smooth spine was obtained by exponentially smoothing the raw spine. **c.** Four additional landmark points, `neck_top, neck`, and `neck_down` (blue), and the contour `centroid` (grey), were calculated.

**Figure 1–Figure supplement 4.**
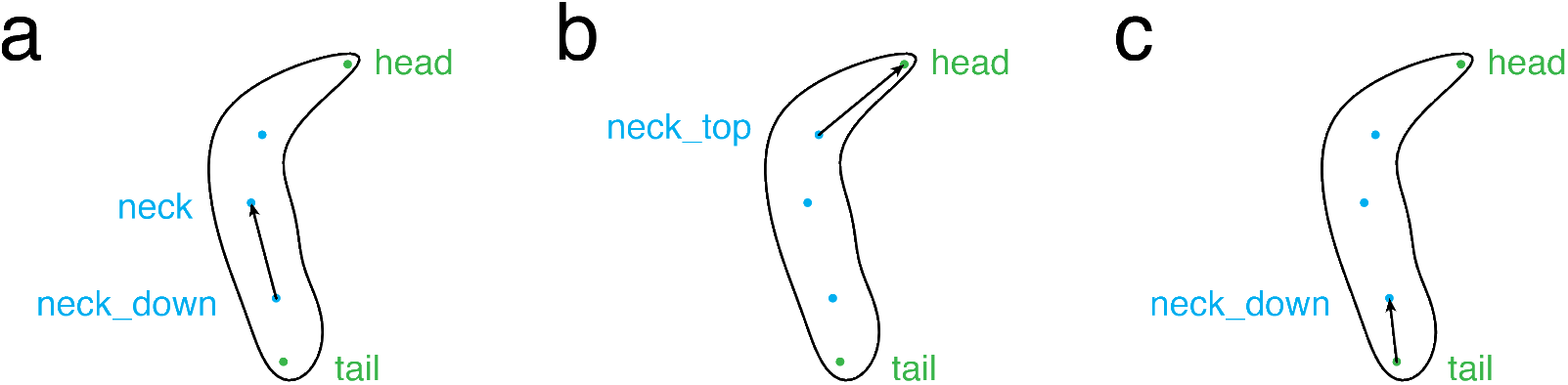
Calculating directionvectors. Three direction vectors were calculated based on `head, tail`, and the landmark points. **a.** `direction_vector` was the normalised vector from `neck_down` to `neck`. **b.** `direction_head_vector` was the normalised vector from `neck_-top` to `head`. **c.** `direction_tail_vector` was the normalised vector from `tail to neck_down`.

**Figure 1–Figure supplement 5.**
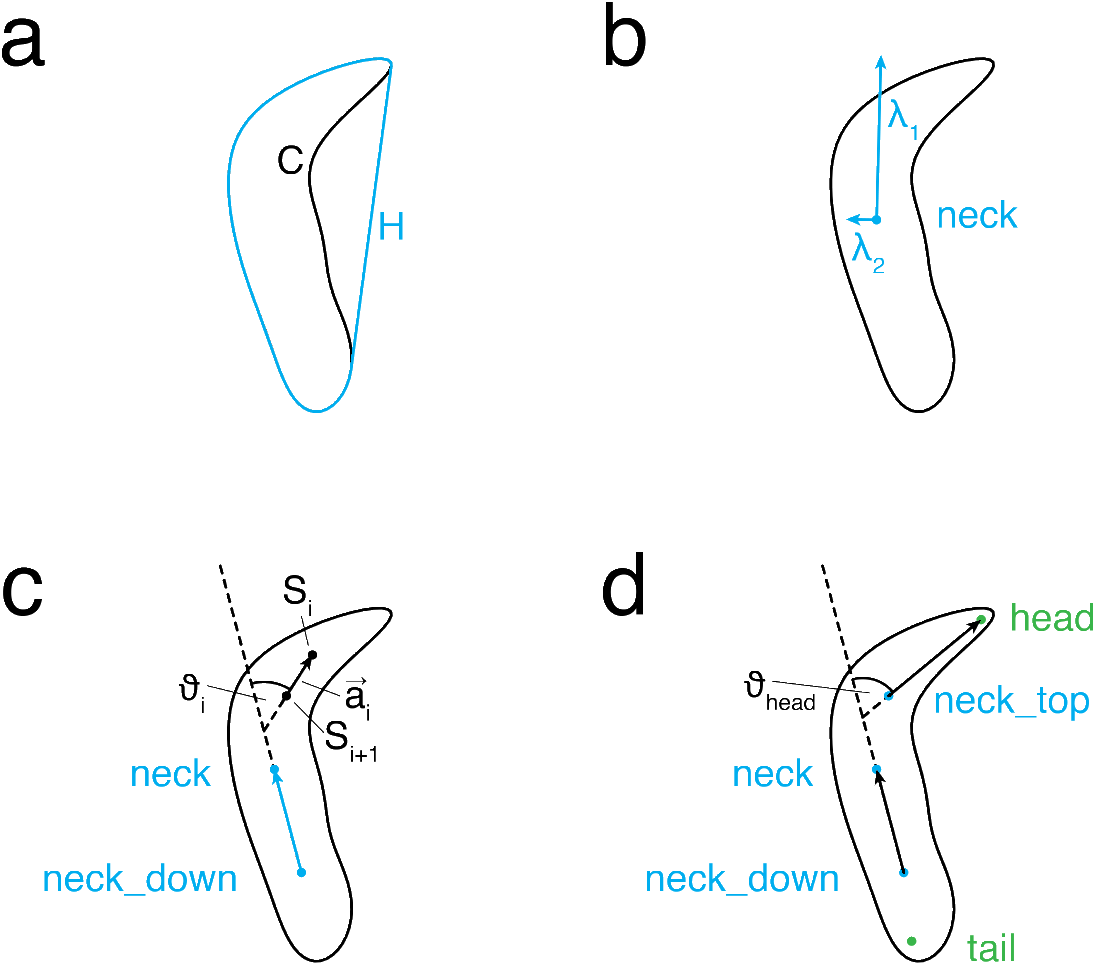
Features describing body shape. **a.** Outline of a larva with contour ***C*** (black) and its convex hull ***H*** (blue). **b.** Shown here are the eigen vectors (blue) of the larval contour (black) structure tensor with respect to `neck` and their corresponding eigen values *λ*_1_ and *λ*_2_. **c.** *ϑ*_*i*_ was defined as the angle between `direction_vector` (blue) and the vector 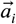 that passed through spine points ***S***_i_ and ***S***_i+1_ (black). **d.** *ϑ*_head_ was defined as the angle between `direction_vector` and `direction_head_vector. head` and `tail` are shown in green.

**Figure 1–Figure supplement 6.**
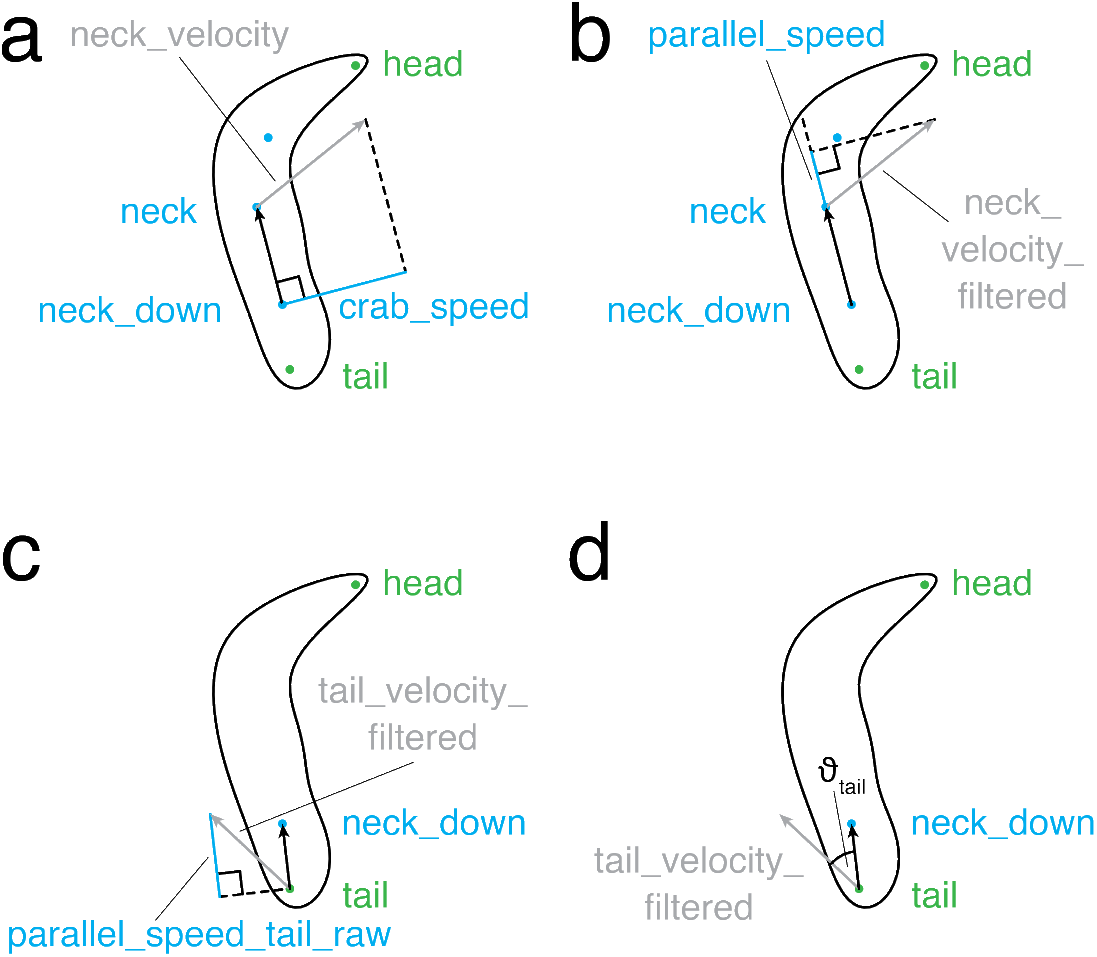
Velocityfeatures. The larval contour is shown in black while head and tail are shown in green. **a.** `crab_speed` (blue) was defined as the component of `neck_-speed` (grey) that was orthogonal to `direction_vector_filtered` (black). **b.** `parallel_speed` (blue) was defined as the component of `neck_speed_filtered` (grey) that was parallel to `direction_vector_filtered` (black). **c.** `parallel_speed_tail_raw` (blue) was defined as the component of `tail_-speed_filtered` (grey) that was parallel to `direction_tail_vector_filtered` (black). **d.** *ϑ*_tail_ was defined as the angle between `tail_speed_filtered` (grey) and `direction_tail_vector_filtered` (black).

**Figure 1–Figure supplement 7.**
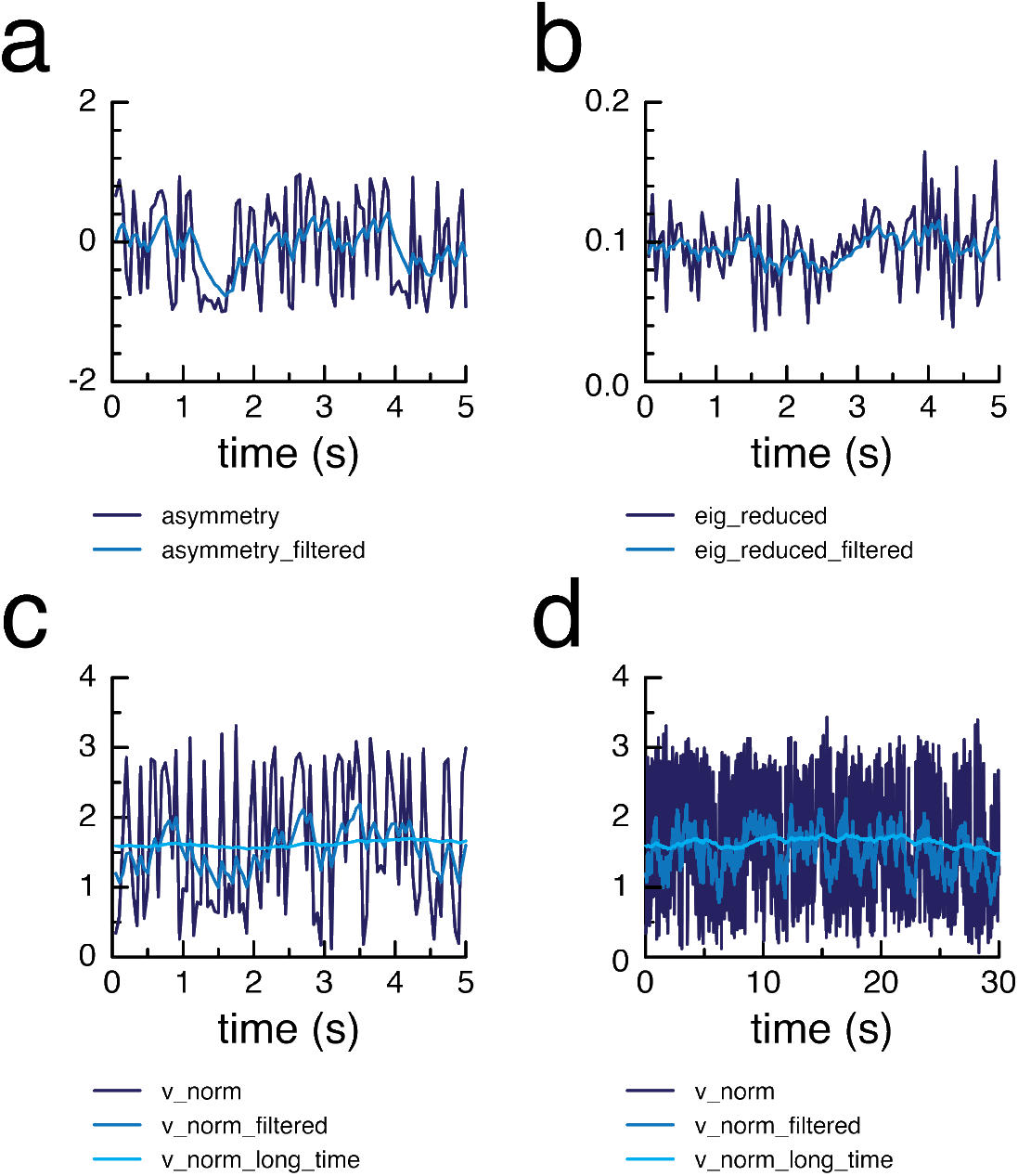
Temporal smoothing of features. **a-b.** Example graphs of raw (dark blue) and filtered (mid blue) `asymmetry` (**a**) and `eig_reduced` (**b**) values over time. **c–d.** Example graphs of raw (dark blue), filtered (mid blue), and long-time filtered (light blue) `v_norm` values over a short (**c**) and a long (**d**) period of time.

**Figure 1–Figure supplement 8.**
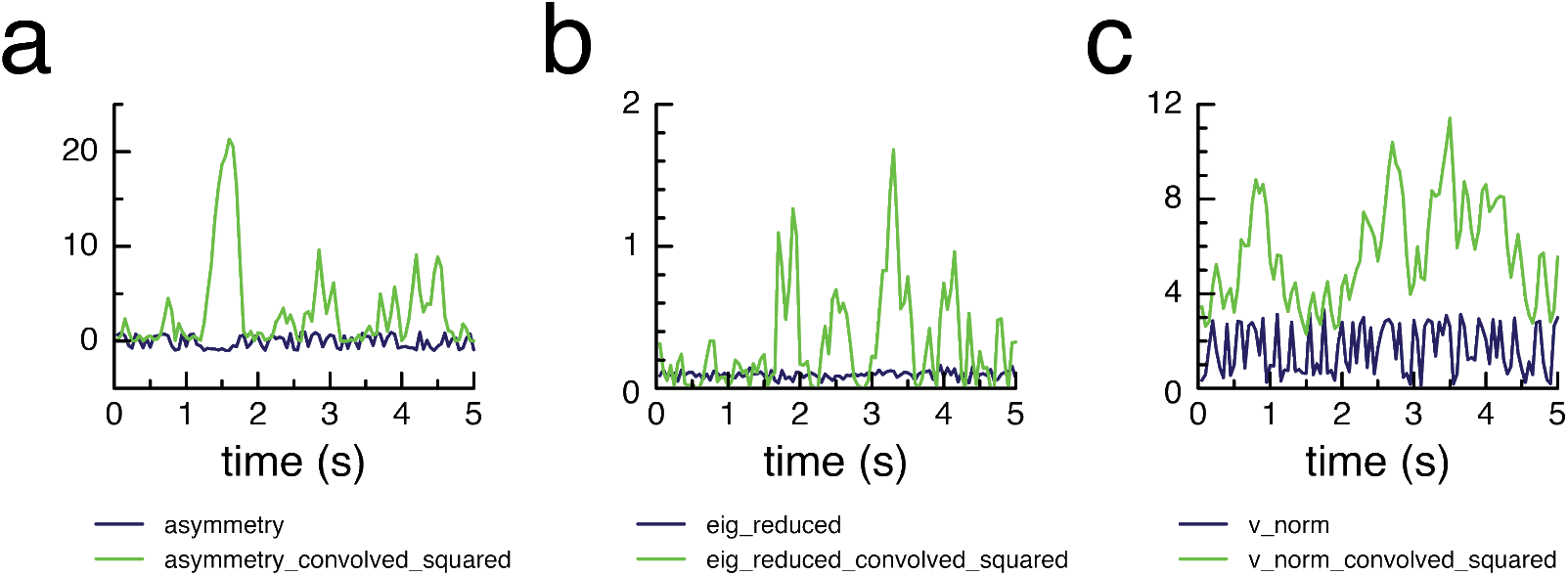
Differentiation by convolution. Example graphs of raw (dark blue) and convolved squared (green) `asymmetry` (**a**), `eig_reduced` (**b**) and `v_norm` (**c**) values over time.

**Figure 3–Figure supplement 1.**
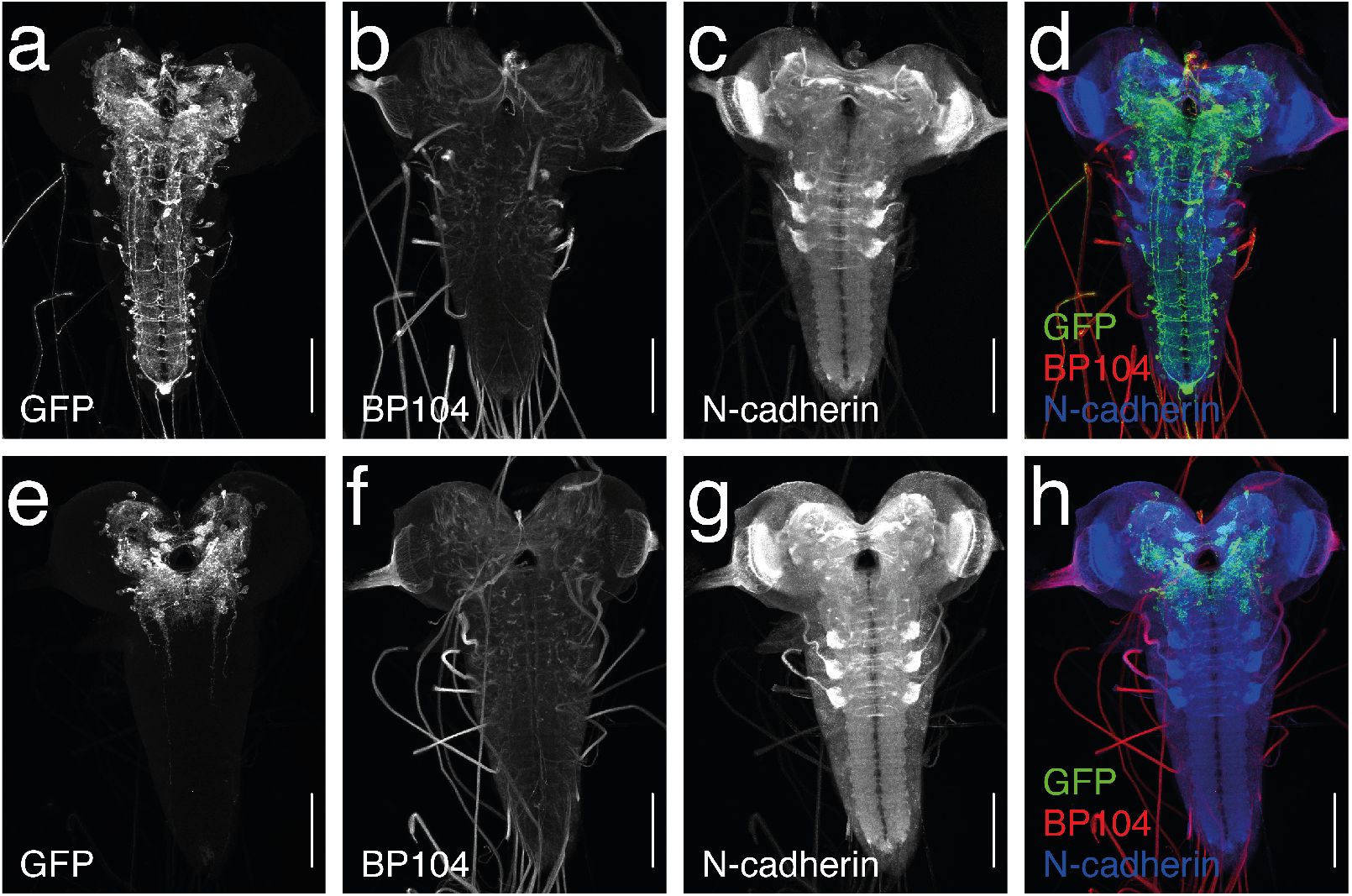
*Ddc-Gal4* expression pattern without and with *tsh-Gal80* restriction. Maximum intensity projections of confocal images obtained after immunohistochemical staining. **a**, **e**; green in **d** and **h.** Targeting a green fluorescent protein (GFP) antibody to the *mVenus* tag of *CsChrimson*. **b**, **f**; red in **d** and **h**. Staining against BP104. **c**, **g**; blue in **d** and **h.** Staining against N-cadherin. **a–d.** *Ddc-Gal4 x UAS-CsChrimson* larvae. Manually counting the cell bodies in the image stacks revealed more than 200 GFP-positive neurons located in thebrain, subesophageal zone (SEZ), and ventral nerve cord (VNC), including the PAM cluster dopaminergic neurons innervating the mushroom body (*n* = 2). This confirmed that *Ddc-Gal4* drives broad expression across the central nervous system (CNS) (*Lundell and Hirsh, 1994*; *Li et al., 2000*). **e–h.** *Ddc-Gal4 x UAS-CsChrimson; tsh-LexA, LexAop-Gal80* larvae. As expected, no GFP-positive neurons were found in the VNC (*n* = 6). *Ddc-Gal4* brain and SEZ expression remained largely unaffected by GAL80, as the GFP-positive neurons in both areas that could be consistently identified in *Ddc-Gal4 x UAS-CsChrimson* larvae (*n* = 3) were also present in *Ddc-Gal4 x UAS-CsChrimson; tsh-LexA, LexAop-Gal80* larvae (*n* = 3). **a–h.** Plan-Apochromat 20x objective, resolution: 592 × 800 pixels, scale bar: 100 μm. Images courtesy of the HHMI Janelia FlyLight team.

**Figure 4–Figure supplement 1.**
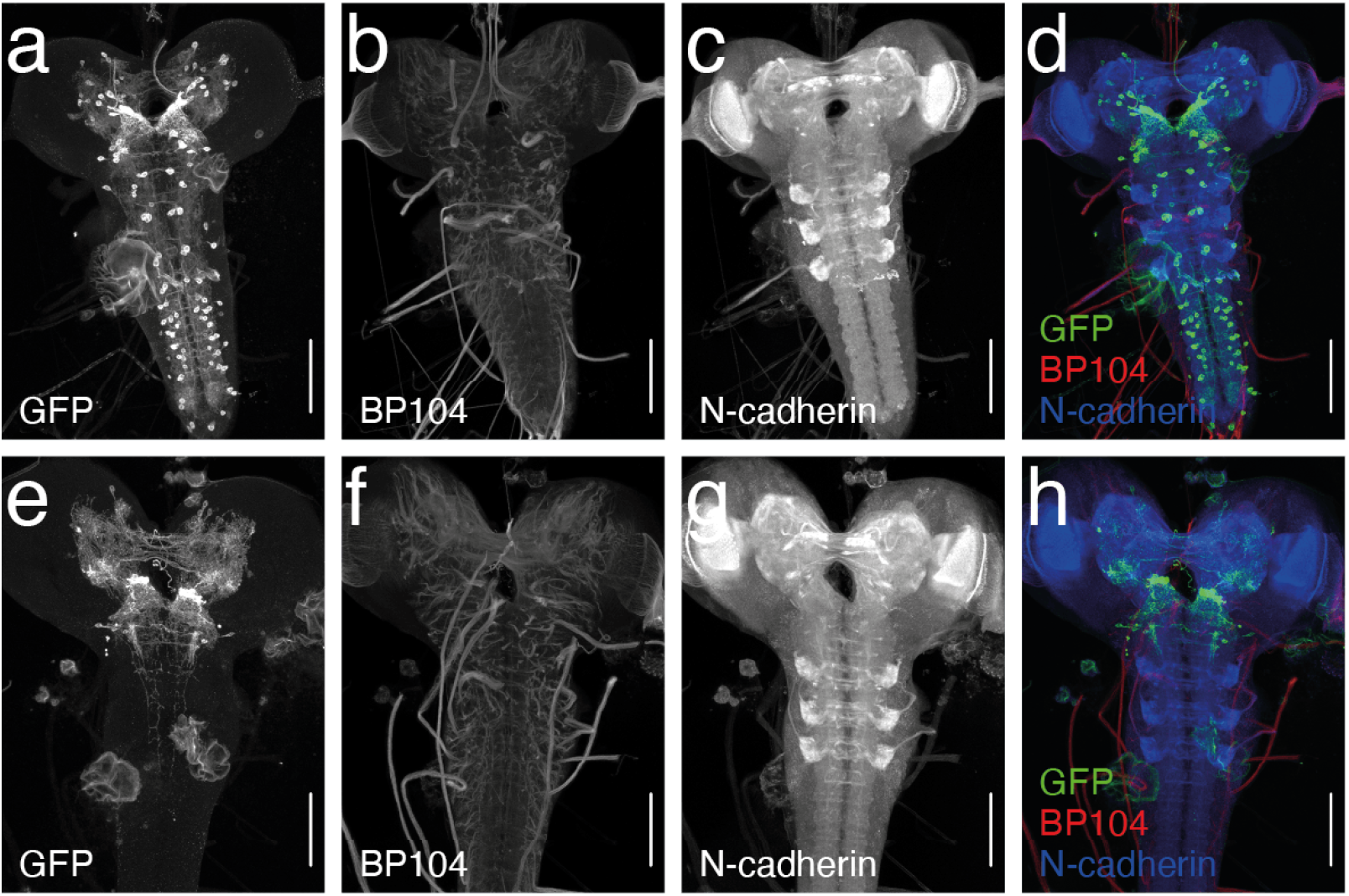
*Tph-Gal4* expression pattern without and with *tsh-Gal80* restriction. Maximum intensity projections of confocal images obtained after immunohistochemistry. **a–d.** *Tph-Gal4 x UAS-CsChrimson* larvae, **e–h.** *Tph-Gal4 x UAS-CsChrimson; tsh-LexA, LexAop-Gal80* larvae. **a**, **e**; green in **d** and **h.** Staining against green fluorescent protein (GFP) antibody targeting the *mVenus* tag of *CsChrimson*. **b**, **f**; red in **d** and **h.** Staining against BP104. **c**, **g**; blue in **d** and **h.** Staining against N-cadherin. **a–h.** Plan-Apochromat 20x objective, resolution: 592 × 800 pixels, scale bar: 100 μm. Image courtesy of the HHMI Janelia FlyLight team.

**Figure 4–Figure supplement 2.**
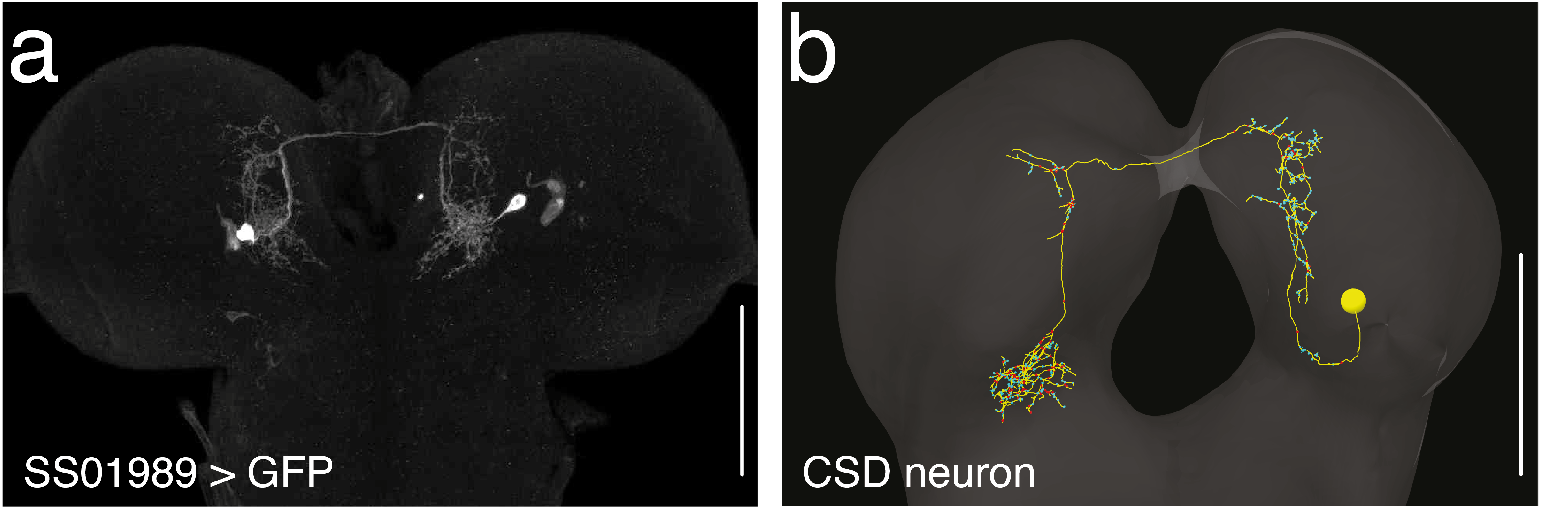
SS01989 exclusively drives expression in the CSD neuron. **a.** Confocal image of a third-instar *SS01989 x UAS-GFP* larva CNS, derived from maximum intensity projections, obtained after immunohistochemical staining against *GFP*. C-Apochromat 40x objective, resolution: 975 × 651 pixels, scale bar: 100 μm. Image courtesy of the HHMI Janelia FlyLight team. **b.** Electron microscopy reconstruction of the contralaterally projecting serotonin-immunoreactive deutocerebral (CSD) neuron from the central nervous system of a first-instar larva (***Berck et al., 2016)***, scale bar: 50 μm.

**Figure 4–Figure supplement 3.**
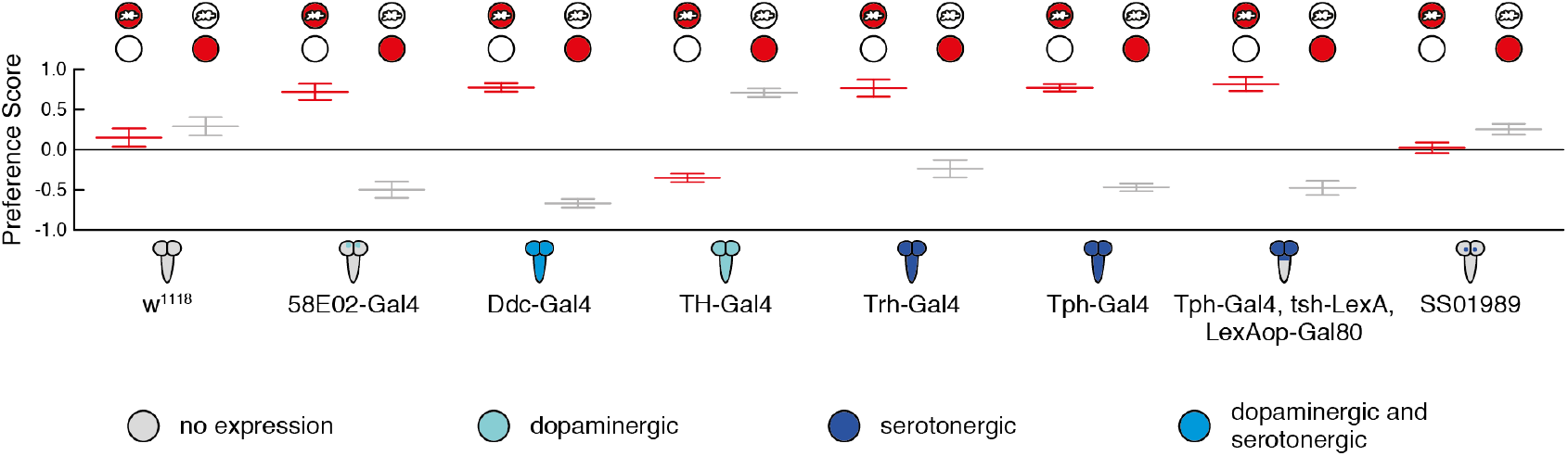
Paired and unpaired group data for olfactory conditioning experiments. The data shown here underlies the performance indices depicted in **Figure 4B**. Gal4 expression depicted as color-coded central nervous system. Preference scores for paired (light/odour, dark/air) and unpaired (dark/odour, light/air) groups are shown in red and grey, respectively.

